# *Camellia sinensis* solvent extract confers trophocidal and cysticidal effects against *Acanthamoeba castellanii*

**DOI:** 10.1101/2022.09.08.507025

**Authors:** Lenu B. Fakae, Mohammad S. R. Harun, Darren Shu Jeng Ting, Harminder S. Dua, Gareth W.V. Cave, Xing-Quan Zhu, Carl W. Stevenson, Hany M. Elsheikha

**Affiliations:** School of Veterinary Medicine and Science, University of Nottingham, Loughborough, LE12 5RD, UK; School of Biosciences, University of Nottingham, Loughborough, LE12 5RD, UK; Rivers State University, Nkpolu - Oroworukwo P.M.B 5080, Port Harcourt, Rivers State, Nigeria; Department of Biomedical Science, Advanced Medical and Dental Institute, Universiti Sains Malaysia, Bertam Campus, Kepala Batas, 13200, Pulau Pinang, Malaysia; Academic Ophthalmology, School of Medicine, University of Nottingham, Nottingham, NG7 2RD, UK; Department of Ophthalmology, Queen’s Medical Centre, Nottingham NG7 2UH, UK; School of Science and Technology, Nottingham Trent University, Nottingham NG11 8NS, UK; Laboratory of Parasitic Diseases, College of Veterinary Medicine, Shanxi Agricultural University, Taigu, Shanxi Province 030801, People’s Republic of China

**Keywords:** *Acanthamoeba castellanii*, *Acanthamoeba* keratitis, *Camellia sinensis*, amoebicidal effect

## Abstract

**Aim:** We examined the anti-acanthamoebic efficacy of solvent extract of *C. sinensis*) and its chemical constituents against trophozoites and cysts of *A. castellanii*.

**Materials and methods:** The effects of *C. sinensis* solvent extract on *A. castellanii* was investigated by using anti-trophozoite, anti-encystation, and anti-excystation assays. The solvent extract was also fractionated using Gas Chromatography and the chemical constituents of *C. sinensis* were tested, individually or combined, against the trophozoites.

**Results:** Trophozoite replication was inhibited within 24-72 h with exposure to 625-5000 µg/mL of *C. sinensis* solvent extract. *C. sinensis* also exhibited a dose-dependent inhibition of encystation, with a marked cysticidal activity at 2500-5000 µg/mL concentrations. Two constituents of *C. sinensis*, namely epigallocatechin-3-gallate and caffeine, significantly inhibited trophozoite replication and encystation at 100 μM and 200 μM, respectively. Cytotoxicity analysis showed that 156.25*-*2500 µg/mL of solvent extract was not toxic to human corneal epithelial cells, while up to 625 µg/mL was not toxic to Madin-Darby Canine Kidney cells.

**Conclusions:** This study shows the anti-acanthamoebic potential of *C. sinensis* solvent extract against trophozoites and cysts. Further pre-clinical studies are required to elucidate the in vivo efficacy and safety of C. *sinensis* solvent extract.

**Graphical abstract:** 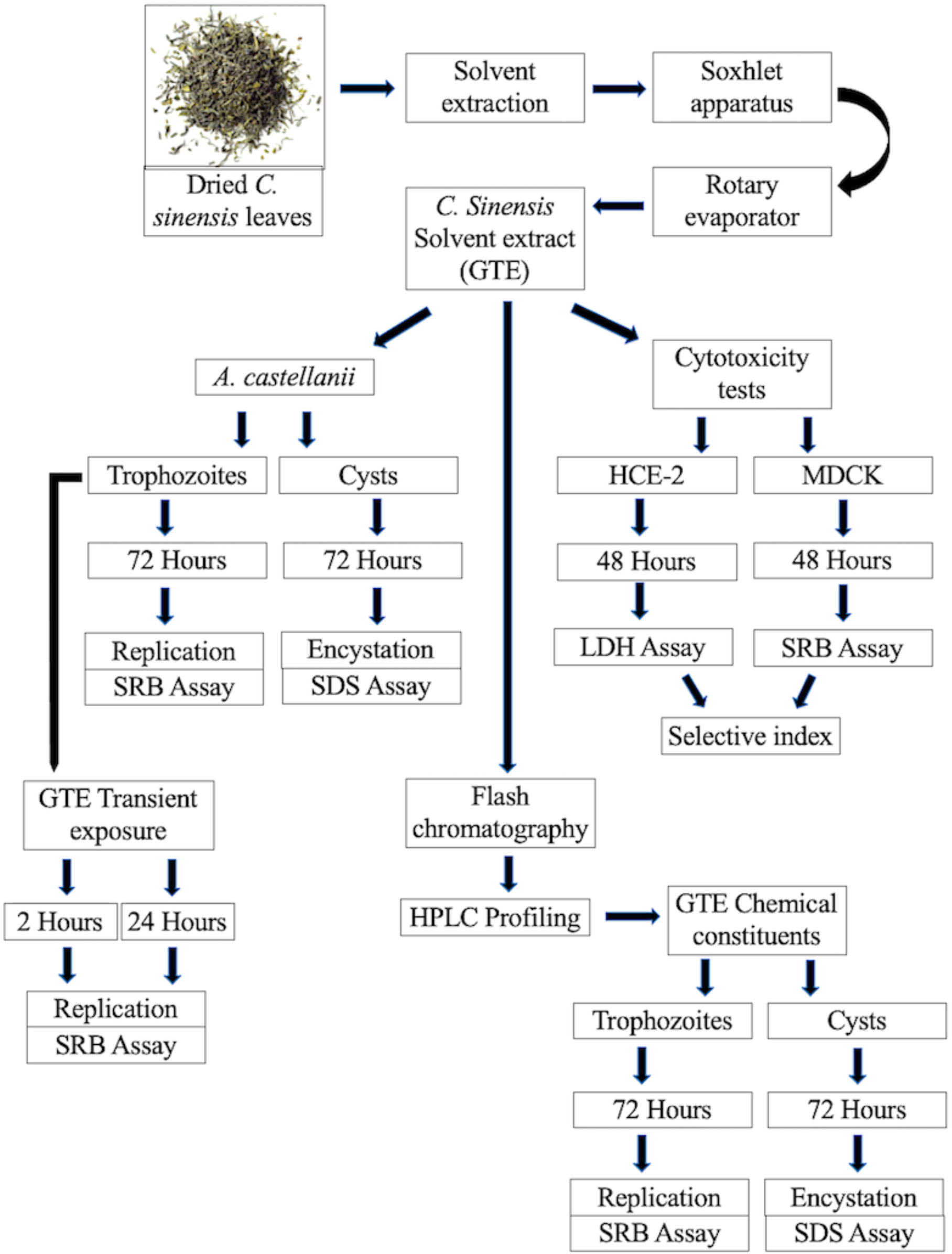

## 1. Introduction

*Acanthamoeba castellanii* is an important parasitic agent that is capable of causing a wide range of serious health conditions, including granulomatous amoebic encephalitis (Kofman and Guarner, 2021), Acanthamoeba keratitis (AK) (Ting et al., 2021a; Ting et al., 2021ba; Carnt et al., 2020), and inflammation of the lungs and skin (Kofman and Guarner, 2021). Polyhexamethylene biguanide (PHMB) and chlorhexidine (Chx) digluconate are the treatment of choice for *Acanthamoeba* infections (Martín-Navarro et al., 2008; Hadas et al., 2017a). Because of the inadequate therapeutic effects of monotherapy with PHMB or Chx, their usage has involved combination with diamidines; propamidine isethionate or hexamidine (Hay et al., 1994; Martín-Navarro et al., 2008; Dart et al., 2009). However, the risks of treatment failure and relapse are high (Siddiqui et al., 2016). Antimicrobial combinations have also been used (Aichelburg et al., 2008; Singhal et al., 2001; Walia et al., 2007; Visvesvara et al., 2007; Garajova et al., 2014), but with limited therapeutic effect.

Alternative therapeutics have been suggested such as photodynamic chemotherapy (by employing the use of light-sensitive medication and a light source to destroy *A. castellanii*) (Ferro et al., 2006), therapeutic corneal cross-linking [by combining ultraviolet light A and riboflavin (B2)] (Khan et al., 2011; Ting et al., 2021c, Said et al., 2021), amniotic membrane transplantation (Ting et al., 2021d), silencing messenger RNA using small interfering RNA (Lorenzo-Morales et al., 2010), and combining nanoparticles with anti-*A. castellanii* drugs for targeted drug delivery to infection sites (Anwar et al., 2018). In the face of therapeutic inadequacies and potential toxicity, researchers are delving into alternate sources of possible treatment using natural sources which have shown anti-acanthamoebic activity (Dodangeh et al., 2018; Hadas et al., 2017a; Hadas et al., 2017b; Saoudi et al., 2017; Mahboob et al., 2017; Kaynak et al., 2018; Sifaoui et al., 2017; Kaya et al., 2019).

Green tea (*Camellia sinensis*), popularly used as a traditional medicine, in the form of infusion, for many purposes including management of diabetes (Mohabbulla Mohib et al., 2016) and hyperglycaemia (Barkaoui et al., 2017). Due to its antioxidant Properties, green or black tea extracts can slow down the progression of eye lens cataract (Thiagarajan et al. 2001). We recently showed the strong anti-acanthamoebic activity of green tea (*Camellia sinensis*; *C. sinensis*) brews against the growth of *A. castellanii* trophozoites and the encystation ability of trophozoites (Fakae et al., 2020). We have also demonstrated the inhibitory effect of (−)-epigallocatechin-3-gallate (EGCG), a major component of *C. sinensis*, on *A. castellanii* growth (Dickson et al., 2020). In addition to EGCG, *C. sinensis* contains catechin, (−)-epicatechin (EC), (−)-epicatechin-3-gallate (ECG), and (−)-epigallocatechin (EGC) (Mukhtar and Ahmad, 2000), and caffeine (Perva-Uzunalic et al., 2006), among other components. Catechins are flavonoids with antioxidant and antiviral capability, which is useful for prevention of many diseases (Sanlier et al., 2018; Yang et al., 2019; Katada et al., 2019; Srichairatanakool et al., 2006; Liou, 2021).

We hypothesised that solvent extraction might yield higher concentration of phytochemicals of C. sinensis and better efficacies compared with the *C. sinensis* brews. The objectives of the present study were to investigate the activity of the solvent extract of *C. sinensis* and its main chemical ingredients against the trophozoites and cysts of *A. castellanii* and evaluate its cytotoxicity against human corneal cell lines and Madin-Darby canine kidney cells.

## 2. Materials and methods

### 2.1. Parasite culture

As described previously (Fakae et al., 2020), *A. castellanii* strain of T4 genotype (American Type Culture Collection; ATCC 30011) was used and maintained in peptone-yeast-glucose (PYG) medium [consisted of proteose-peptone 0.75% (w/v), yeast extract 0.75% (w/v) and glucose 1.5% (w/v)] in T75 Nunclon^®^ tissue culture flasks (ThermoFisher Scientific, Loughborough, UK) at 25°C in a humidified Stuart™ SI30H Hybridization bench top oven/shaker (Cole-Parmer Ltd, Staffordshire, UK) without rocking. The culture medium was refreshed ∼15 h before the commencement of each experiment to promote the growth of active trophozoites.

### 2.2. Solvent extraction of C. sinensis

The solvent extraction of *C. sinensis* was performed using a 2-step procedure with Soxhlet apparatus. Methanol and acetonitrile were used in step 1 and step 2, respectively. Dried *C. sinensis* was ground to powder in a laboratory mill and weighed into in large sized Whatman^®^ cellulose extraction thimbles (Sigma-Aldrich, UK) with 10 μm nominal particle retention, which were plugged with cotton wool and placed in Soxhlet apparatus. 500 mL of the solvent was poured through the thimble into a 1000 mL round bottom flask allowing for an initial extraction of the sample prior to the commencement of the extraction cycles. The Soxhlet apparatus was placed on a hotplate (Radley’s Tech, Germany) and the temperature was set to 70°C for methanol and 85°C for acetonitrile, respectively, and the solvent extraction was stirred by using a magnetic stirrer at 25xg. The solvent extraction was carried out for 48 h; 24 h for each step. Both solvent extracts were pooled together and after cooling the suspensions were concentrated to dry matter using a vacuum with rotary evaporator (Rotavapor® R-300,BUCHI, Flawil, Switzerland) and water bath set to 40°C.

### 2.3. Preparation of solvent extract solutions for testing

*C. sinensis* solvent extract solution was prepared at a stock concentration of 10,000 µg/mL containing 0.025% DMSO. Firstly, 500 mg of the sample was weighed into a 50 mL falcon tube and 2.5 mL of dimethyl sulfoxide (DMSO) was added. This was placed in an ultrasonicator bath (Elma, Germany) for 15 min to form a homogeneous solution. Then, 47.5 mL of PGY media (which was used throughout the experiments as a negative control) was added to the initial solution to make up 50 mL of stock solution. The solution was filtered sterile using a 0.22 µL Millipore syringe filter (MILLEX^®^-HA brand, Ireland). Two-fold serial dilutions of *C. sinensis* solvent extraction solution (from 5000 µg/mL to 156.25 µg/mL) were prepared from the stock solution using PGY as a diluent. In view of the possible cytotoxic effect of DMSO, which could erroneously increase sample anti-amoebic activity, respective concentrations of DMSO were used as vehicle controls and the effects were accounted for during the final calculation of anti-acanthamoebic activity. For the cytotoxicity assays, *C. sinensis* serial dilutions (v/v) were prepared as described above using the respective cell culture medium as the diluent.

For *C. sinensis* encystation medium, a stock concentration of 10,000 µg/mL was achieved by initially preparing a solvent extract solution as described above but using distilled water instead of PGY media. For the hyperosmotic encystation media, 10 g of glucose monohydrate, 0.48 g of magnesium chloride (Sigma-Aldrich) and one phosphate-buffered saline (PBS) tablet (Gibco^®^, Life technologies, ThermoFisher Scientific, UK) was dissolved in 100 mL of the distilled water-*C. sinensis* solvent extract solution. This was stirred on a magnetic stirrer for 30 min and the suspension filtered into a sterile bottle using a 0.45 µL Millipore syringe filter (MILLEX^®^-HA, Ireland). The standardized encystation medium (negative control) was prepared by using distilled water as a solvent. The positive control included standardized encystation medium supplemented with 5% PMSF solution (Sigma life science, Switzerland).

### 2.4. Cytotoxicity tests

Established lactate dehydrogenase (LDH) assay and sulforhodamine B (SRB) assay were used to measure the cytotoxicity of *C. sinensis* solvent extract against HCE-2 cells and Madin-Darby Canine Kidney (MDCK) cells, respectively. Both experiments were conducted in technical triplicate as three independent experiments.

HCE-2 cells (CRL11135, ATCC, Manassas, Virginia, USA) were seeded into a Thermo Scientific™ Nunc MicroWell 96-well plate (ThermoFisher Scientific, Loughborough, UK), at 7.5×10^3^ cells/well and grew to 80-90% confluency, in the presence of growth media (consisting of keratinocyte serum free medium supplemented with human recombinant epidermal growth factor, bovine pituitary extract, hydrocortisone, and insulin) (Bagshaw et al., 2020). HCE-2 cells were subsequently incubated with 1:2 serial concentration of *C. sinensis* starting from 5000 µg/mL to 156.3 µg/mL) for 48 h. The culture medium was used as a negative control. In view of the potential sedimentation of *C. sinensis* (which could erroneously increase the optical density (OD) reading, hence toxicity), additional wells containing the same 1:2 serial concentration of *C. sinensis* (5000-156.3 µg/mL), but without HCE-2 cells, were included so that any effect of precipitated *C. sinensis* on the OD reading can be determined and accounted for during the calculation of cytotoxicity. The LDH assay (ThermoFisher Scientific, UK) was performed as per the manufacturer’s instructions. At 24 h and 48 h post-treatment, 50 µl of supernatant was obtained from each well and OD_490-680_ was measured using BMG Clariostar microplate reader (BMG LABTECH Ltd., Aylesbury, United Kingdom). Cytotoxicity (%) was calculated using the following formula: [((_ltreatment_ – _INC_) / (_IPC_ – _INC_)) x 100; I = intensity].

MDCK cells were seeded in 96-well microplates at a density of 5×10^3^ cells/well in 100 µl Gibco Dulbecco’s Modified Eagle Medium (DMEM) (Gibco^®^, Life technologies, ThermoFisher Scientific, UK). After 48 h of incubation in a humidified atmosphere of 5% CO_2_ at 37°C, the medium was removed using an electronic pipette (Gilson, France), and the cultures were treated with the solvent extract at the same above-mentioned concentrations and incubated for 3, 24, and 48 h. Parallel negative control wells contained MDCK cells with the respective medium only, without *C. sinensis*. Positive control included wells treated with 3% SDS. At each time point of incubation, cell proliferation was determined using the SRB assay as described previously (Ortega-Rivas et al., 2016).

### 2.5. Evaluation of the trophocidal activity

We investigated the dose-dependent effect of *C. sinensis* solvent extract on the growth kinetics of trophozoites. Trophozoites (3.2×10^5^) were treated with solvent extract at concentrations of 5000, 2500, 1250, and 625 µg/mL, and seeded into T25 Thermo Scientific™ Nunc tissue culture flasks (ThermoFisher Scientific, Loughborough, UK), which were incubated in a humidified Stuart™ SI30H Hybridization bench top oven/shaker at 25°C. Controls included an equal number of trophozoites in PYG only (negative control) or in PYG supplemented with 0.02% chlorhexidine (Chx) (positive control). After 24, 48 and 72 h, trophozoites were counted using a hemocytometer (Neubauer-improved bright line, Marienfeld, Germany) and a CETI inverted microscope (Medline Scientific, UK) to determine the effect of each concentration on the growth rate of the trophozoites. All experiments were performed in technical triplicate as three independent experiments.

### 2.6. Selectivity index

The *C. sinensis* solvent extract concentration that caused 50% inhibition of mammalian cell growth was expressed as 50% cytotoxic concentration (CC_50_). The CC_50_ of *C. sinensis* on HCE-2s and MDCKs were calculated by plotting dose-response curves followed by simple linear regression analysis using Graph Pad Prism 7 software. The calculation of half-maximal inhibitory concentration (IC_50_), which is the concentration of *C. sinensis* solvent extract that caused a 50% reduction in the growth of *A. castellanii* trophozoites compared to the control, was performed using Graph Pad Prism 7’s nonlinear regression (curve fit) built-in analysis tool of dose-response inhibition (three parameters). The selectivity index (SI), which represents the ratio of the CC_50_ for mammalian cells to the IC_50_ for *A. castellanii* was calculated by comparing the cytotoxicity of *C. sinensis* for HCE-2 cells and MDCK cells to that of trophozoites.

### 2.7. Effects of transient exposure on trophozoites

3×10^5^ trophozoites were seeded in 15 mL falcon tubes treated with 5 mL of solvent extract in concentrations of 5000, 2500, 1250, and 625 µg/mL, with PYG media as a negative control and 0.02% CHX in PYG as a positive control. After 2 and 24 h post-treatment, trophozoites were harvested, washed twice with PYG medium to remove traces of *C. sinensis*. Then, trophozoites were suspended in 5 mL of fresh PYG, seeded in T25 tissue culture flasks, and further incubated for 24, 48 and 72 h. At each of these incubation time points, the number of trophozoites was counted using a hemocytometer to determine the growth inhibitory effect of the treatment on trophozoites. The treated trophozoites were also examined optically using a Leica DMIL CMS (Germany) inverted microscope with a Leica Application Suite (LAS version 4.3) to identify any morphological alterations caused by exposure to *C. sinensis*.

### 2.8. Inhibitory effect of C. sinensis on cyst formation

6×10^5^ trophozoites were treated with *C. sinensis* solvent extract hyperosmotic solution in concentrations of 5000, 2500, 1250, and 625 µg/mL, and were seeded in T25 tissue culture flasks. The negative control included trophozoites treated with hyperosmotic encystation medium. The positive control included trophozoites treated with hyperosmotic encystation medium plus 5% PMSF solution. The encystation rate was determined by counting the number of trophozoites versus cysts for each concentration after 24, 48 and 72 h of treatment using a hemocytometer. Also, after 72 h, the cultures were treated with 0.5% sodium dodecyl sulphate (SDS) and left for 60 min at room temperature to digest any remaining trophozoites. Then, pre-, and post-SDS treatment counts were performed and the encystment percentage was determined using the formula:

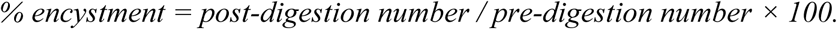

### 2.9. Inhibitory effect of C. sinensis on excystation

2 × 10^5^ *A. castellanii* cysts were treated with *C. sinensis* solvent extract in concentrations of 5000, 2500, 1250, and 625 µg/mL, and the suspensions were seeded in 15 ml falcon tubes. The suspensions were mixed using a vortex mixer (Thermo Scientific, Colchester, United Kingdom) to ensure a homogenous mixture. The samples were placed on a rack incubated in a Stuart oven at 25°C. Negative control included cysts in PYG only, while the positive control included cysts in PYG supplemented with 0.02% CHX. After incubation for 24, 48 and 72 h, the culture was counted to determine the cyst to trophozoite excystation ratio using a hemocytometer and a CETI inverted microscope. The treated cysts were also examined optically using a Leica DMIL CMS (Germany) inverted microscope with a Leica Application Suite (LAS version 4.3) to identify any morphological alterations during excystation caused by exposure to *C. sinensis* solvent extract.

### 2.10. Scanning electron microscopy (SEM)

*A. castellanii* trophozoites and cysts treated with solvent extract in respective experiments were harvested, fixed, and dehydrated using a graded series of ethanol. After dehydration, critical point drying was performed by infiltrating the samples in hexamethyldisilazane (Acros, Germany) for 5 min twice to further enhance drying of the samples. The samples were mounted on aluminium SEM stubs with a double-sided carbon sticker and sputter coated with gold prior to SEM imaging. Images were obtained with varying magnifications using a JEOL JSM-7100F LV Scanning Electron Microscope (JOEL, Tokyo, Japan) operating at an accelerating voltage of 10 kV and a working distance of 10 mm. Data was collected using the lower secondary electron detector.

### 2.11. Partitioning of C. sinensis solvent extract using flash chromatography

After the solvent extraction of *C. sinensis*, a sample was taken out for flash column chromatography using Puriflash 5.125Plus (Interchim, Montluçon, France) to identify the fractions in *C. sinensis* solvent extract. To identify which method separates more fractions, two runs were conducted with different combinations of solvents as mobile phases. One mobile phase (A and B) included 0.1% formic acid in water and water + 0.1% formic acid in methanol, while the second (A and B) included 0.2 % acetic acid in water + acetonitrile at a 91:9 ratio and water + acetonitrile at a 20:80 ratio. Rather than using a single HCC18 Flash Column, two flash columns were used; puriflash C18HC spherical silica flash II column with particle size 50 µm for high capacity weighing 40 g, and puriflash C18HP spherical silica flash II column with particle size 15 *µ*m for high performance weighing 12 g. The injection mode was liquid with 1 ml of *C. sinensis* solvent extract, the detection was 258 nm and 278 nm with a flowrate of 26 ml/min. The pressure was set to 9 bars and auto gradient optimization mode was used. The elution profile for the first mobile phase (solvent B) was 5% to 95% holding gradient while analytes showed its maximum absorbance, over a run time of 60 min. The elution profile for the second mobile phase (solvent B) was auto gradient optimization holding gradient while analytes showed its maximum absorbance over a run time of 59 min and only fractions of monitored peaks were collected.

### 2.12. HPLC profiling of C. sinensis solvent extract

After collecting fractions of monitored peaks observed during flash chromatography of *C. sinensis* solvent extract, the fractions collected were characterized using ultra-high performance liquid chromatography quadrupole time of flight tandem mass spectrometry (UHPLC-QTOF-MS) (Waters, Milford, MA, USA) to identify the analytes present in these fractions. Two solvents (A and B) were used as mobile phase. A was 0.1% formic acid in water + methanol at a 90:10 ratio while B was 0.1% formic acid in methanol. A Restek C18 Raptor™ column (Restek Thames, High Wycombe, United Kingdom) was used. The injection mode was performed by automatic liquid injection with injection volume of 1 mL (50 µl crude solvent extract diluted in 950 µl of LC-MS grade methanol). The flow rate was 0.25 ml/min with a run time of 15 min. The elution gradient of solvent B was 10% ramped to 95% over a period of 15 min while that of solvent A was 95% ramped to 10% over a period of 15 min. The chromatographic data were analyzed, and chemical composition of each component was confirmed by matching the chemical formula with references in the library of MassLynx^®^ software (version 4.1, Waters, Milford, MA, USA).

### 2.13. Inhibitory effects of C. sinensis chemical constituents on trophozoites

The chemical substances of the 10 active components namely theogallin, theobromine, epigallocatechin (EGC), caffeine, epicatechin gallate (ECG), epigallocatechin gallate (EGCG), epicatechin (EC), catechin, kaempferol, myricetin, all identified in *C. sinensis* by chromatography and HPLC profiling of the *C. sinensis* solvent extract were purchased from Sigma-Aldrich (UK). Stock solutions of each standard were prepared with PGY to a final concentration of 1,000 µM. Two-fold serial dilutions of each *C. sinensis* chemical standards (from 200 µM to 3.12 µM) were used individually to investigate their individual inhibitory effect on the growth kinetics of trophozoites using the colorimetric SRB assay. For each chemical standard, trophozoites were seeded at 2.5×10^3^ trophozoites/well in 96-well microtiter plates. Each well received 100 µl of the testing chemical standards. After testing each chemical standard individually, we further tested the efficacy of all 10 chemical standards combined. Another experiment warranted us to combine 2 of the most potent chemical standards to determine any potential synergist effect. For each experiment, negative control wells received only 100 µl PYG medium/well, while positive control wells received 100 µl 3% SDS, which was added 30 min before commencement of the SRB assay. The plates were incubated in a Stuart oven at 25°C. Then, at 3, 6, 24, 48 and 72 h post-incubation, the colour absorbance of each well was measured as previously described (Fakae et al., 2020; Ortega-Rivas et al., 2016). All experiments were performed in technical triplicate as three independent experiments.

### 2.14. Inhibitory effect of C. sinensis constituents on encystation

To determine the inhibitory effect of specific constituents of *C. sinensis* on encystation, stock solutions of 1000 µM of EGCG and theobromine, respectively were used as solvent for preparing serial dilutions of hyper osmotic solutions of each chemical constituent for anti-encystation testing. Approximately 6.2×10^5^ *A. castellanii* trophozoites, seeded in T25 tissue culture flasks, were treated with 500, 250, 125, and 62.5 µM of encysting solutions of three *C. sinensis* chemical constituents; EGCG, theobromine and EGCG-theobromine mixed in a 1:1 ratio. The negative and positive controls were the same as described in encystation experiments above. The experiments were performed in triplicate.

### 2.15. Statistical analysis

All statistical analyses were performed using GraphPad Prism Version 9.0.0 (GraphPad Software Inc., CA USA). One-way and two-way analysis of variance (ANOVA) with Turkey’s multiple comparison test were used to compare groups. The results are expressed as mean ± standard error of mean (SEM). *P* < 0.05 was considered statistically significant and *P* < 0.01 was considered as highly statistically significant.

## 3. Results

### 3.1. Cytotoxicity against mammalian cells

The HCE-2 cells were exposed to serial concentrations of solvent extract at doses of 156.25*-*5000 µg/mL (Fig. 1A). By 24 h post-treatment, no significant differences were observed in the percentage toxicity between control and treatment groups at concentrations 156.25*-*1250 µg/mL, indicating that these concentrations were not toxic to HCE-2 cells (*P* > 0.9999). However, 2500 *µ*g/mL and 5000 *µ*g/mL concentrations showed significant increase in the toxicity compared with the control (*P* = 0.0129 and *P* < 0.0001, respectively). At 48 h post-treatment, 156.25-2500 *µ*g/mL showed no significant difference to the control (*P* > 0.9999), indicating that these concentrations were not toxic to HCE-2. However, 5000 µg/mL showed a significant increase in cytotoxicity in HCE-2 compared with the control (Fig. 1A). The cytotoxicity observed in 2500 µg/mL at 24 h could be a transient because toxicity had significantly dropped at 48 h indicating that 2500 µg/mL was not toxic to HCE-2 cells.

**Fig. 1.**
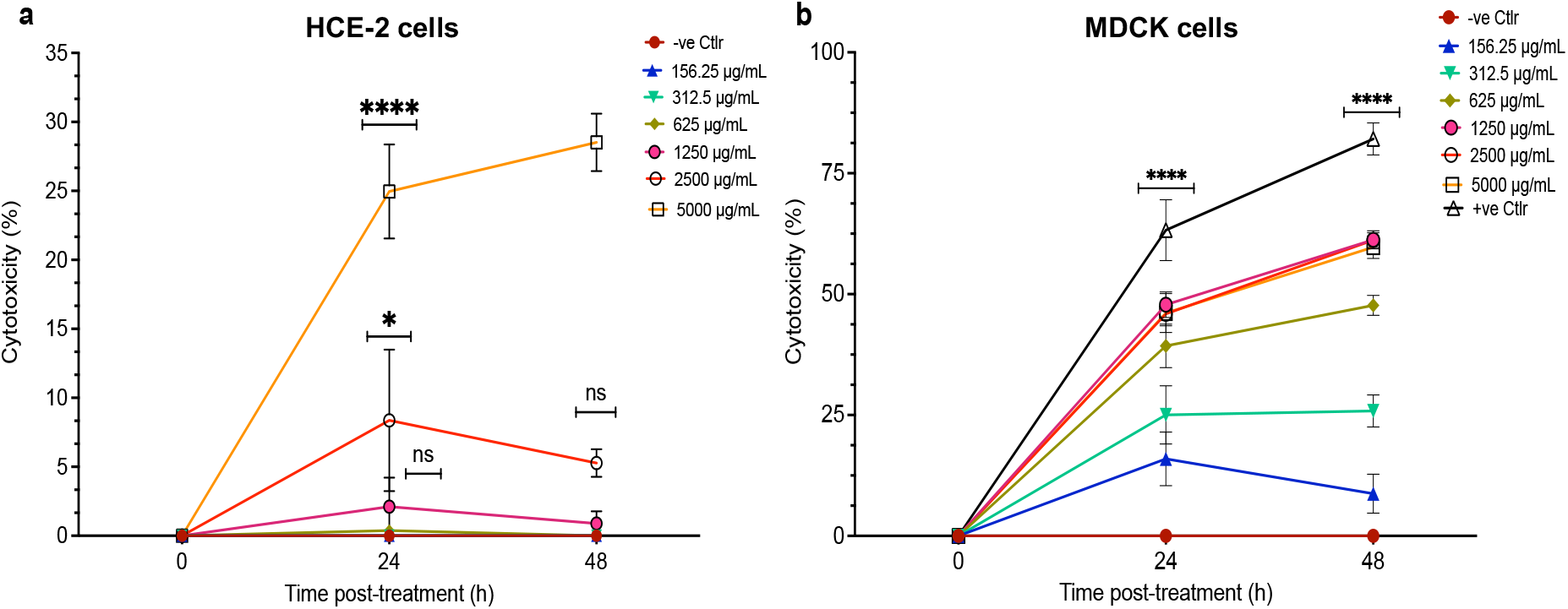
The toxicity of *C. sinensis* solvent extract on cultured human corneal epithelial (HCE-2) cells and Madin-Darby Canine Kidney (MDCK) cells. Cultured cells were exposed to the indicated concentrations and cytotoxicity was investigated using lactate dehydrogenase assay and Sulforhodamine B assay for HCE-2 cells and MDCK cells, respectively. (a) Cytotoxicity against HCE-2 cells showed no significant difference between the control and the concentrations 156.25-2500 µg/mL at 48 h post treatment (*P* > 0.999). However, there was a significant difference between the control and 5000 µg/mL treatment group at 48 h. This indicates that only the highest concentration is toxic to HEC-2 cells, with less than 30% cytotoxicity. (b) Cytotoxicity on MDCK cells showed significant difference between the control (DMEM-treated) and *C. sinensis*-treated groups at 24 h and 48 h post treatment. Despite the difference, concentrations 156.25 µg/mL, 312.5 µg/mL and 625 µg/mL showed an increase in cell replication from 0 h to 48 h, with average percentage cytotoxicity of 10%, 25%, and 45%, respectively at 48 h. There were significant differences between these lower concentrations and higher concentrations 1250 µg/mL, 2500 µg/mL and 5000 µg/mL which recorded an average percentage cytotoxicity of up to 60%. (**** *P* < 0.0001; ns, non-significant)

We determined the toxicity of *C. sinensis* solvent extract to MDCK cells (Fig. 1B). MDCK cells treated with 156.25*-*5000 µg/mL for 24 h showed an increase in cell replication, although the rates were significantly lower than the negative control (*P* < 0.0001), suggesting cytotoxicity caused by *C. sinensis*. At 48 h post-treatment, although MDCK cells treated with various *C. sinensis* concentrations continued to grow, the statistical difference with the control was still significant (*P* < 0.0001).

### 3.2. Inhibitory effects of C. sinensis extract on trophozoites

The inhibitory effect of *C. sinensis* solvent extract on on trophozoites replication was determined using a hemocytometer (Fig. 2A). The two-way ANOVA revealed a significant main effect of time (h) (F (3, 6) = 1148, *P* < 0.0001), concentration (µg/mL) (F (7, 14) = 26766, *P* < 0.0001), and time x concentration interaction (F (21, 42) = 2726, *P* < 0.0001). At 24 h post-treatment, post hoc comparisons of treatment groups with negative control (PGY), vehicle control (PGY+DMSO), and positive control (CHX) showed an increase in the number of trophozoites exposed to 312.5 µg/mL concentration following the trend of the negative control DMSO control (*P* < 0.0001) (Fig. 2A). The other treatment concentrations, 625*-*5000 µg/mL showed a significant decrease in trophozoite numbers compared to the negative control (*P* < 0.0001). At 48 h, trophozoites treated with 312.5 µg/mL continued to grow with no significant difference from the negative and vehicle control groups (*P* = 0.0862). The higher concentrations continued to cause reduction in the number of trophozoites, though still with significant increase from the positive control (*P* < 0.0001). At 72 h post-treatment, 312.5 µg/mL still showed increase in trophozoites, while the other higher concentrations showed decrease in trophozoites with no significant difference between these higher doses. The effect of the two highest does 2500 µg/mL an- 5000 *µ*g/mL was comparable to the effect of positive control as they completely eradicated *A. castellanii* trophozoites (*P* > 0.9999).

**Fig. 2.**
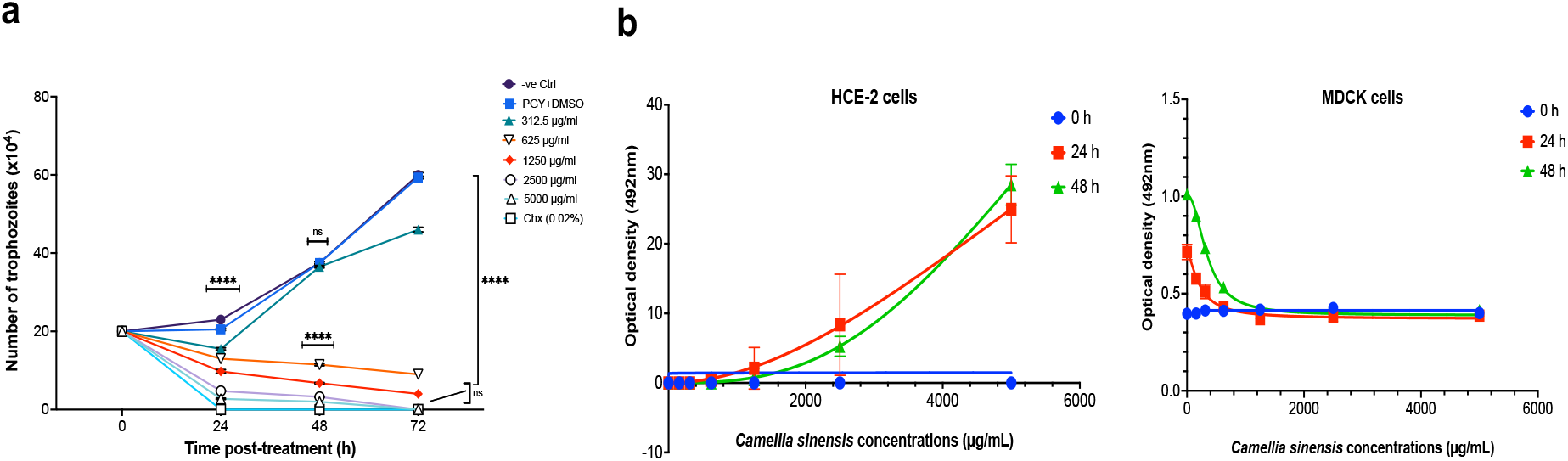
Inhibitory effects and selectivity of *C. sinensis* extract on trophozoites. (a) Growth inhibitory effect of *C. sinensis* on *A. castellanii* trophozoites at 24, 48, and 72h post treatment doses of 312.5-5000 µg/mL. At 24 h, there was a significant decrease in trophozoite number for all *C. sinensis* treatment groups compared with the negative control PGY and vehicle control PGY+DMSO. The decrease was progressive for 625-5000 µg/mL up to 72 h, while 312.5 µg/mL allowed for trophozoite replication. At 48 h, 312.5 µg/mL allowed trophozoites to continue their growth with no significant difference from the negative and vehicle controls (*P* = 0.0862). The higher concentrations continued to reduce the trophozoites’ number, though with significant increase compared with the positive control. At 72 h post-treatment 2500 µg/mL and 5000 µg/mL mimicked the effect of Chx and destroyed *A. castellanii* trophozoites with no statistical difference (*P* > 0.9999). (b) The half-maximal inhibitory concentration (IC_50_) of *C. sinensis* solvent extract for HCE-2 cells and MDCK cells expressed as 50% cytotoxic concentration (CC_50_) was determined to be 8891 ± 1135 µg/mL at 24 h and 6916 ± 199 µg/mL at 48 h for HCE-2 cells and 222.1 ± 25.06 µg/mL at 24 h and 346.3 ± 12.89 µg/mL at 48 h for MDCK cells. (**** *P* < 0.0001; ns, non-significant)

### 3.3. Selectivity index of C. sinensis extract

The half-maximal inhibitory concentration (IC_50_) of *C. sinensis* solvent extract against *A. castellanii* trophozoites was 1309 ± 517.1 µg/mL at 24 h and 409 ± 8.09µg/mL at 48 h (Fig. 2A). The CC_50_ for HCE-2 cells and MDCK cells were 8891 ± 1135 µg/mL at 24 h and 6916 ± 199 µg/mL at 48 h, and 222.1 ± 25.06 µg/mL at 24 h and 346.3 ± 12.89 µg/mL at 48 h, respectively (Fig. 2B). This indicates that *C. sinensis* solvent extract has selective cytotoxicity for *A. castellanii* with a selective index = 6.8 at 24 h and = 16.9 at 48 h, when tested against HCE-2 cells. However, *C. sinensis* solvent extract had a selective index = 0.17 and 0.85, when tested against MDCK cells at 24 h and 48 h, respectively. These results demonstrate the less sensitivity of HCE-2 cells, compared to MDCK cells, to *C. sinensis* solvent extract treatment.

### 3.4. Transient exposure to C. sinensis extract

Transient, 2 h and 24 h, effect of *C. sinensis* solvent extract at 625 µg/mL, 1250 µg/mL, and 5000 µg/mL on trophozoite replication was determined using a hemocytometer (Fig. 3). Post-hoc comparisons for 2 h exposure showed a significant difference (*P* > 0.05) between the parasite growth inhibition caused by CHX and the three concentrations. Likewise, for 24 h exposure the post-hoc comparisons also showed a significant decrease (*P* > 0.05) between the parasite growth inhibition caused by CHX and the three concentrations. *C. sinensis* solvent extracts at 625 µg/mL, 1250 µg/mL, and 5000 µg/mL did not exhibit any sustained trophozoite growth inhibition after a transient exposure of 2 h and 24 h to the trophozoites (Fig. 3). For 2 h and 24 h exposure, despite the significant increase in trophozoite numbers between negative control and *C. sinensis* concentrations (*P* < 0.0001), there was continuous replication of the trophozoites between 24 h to 72 h.

**Fig. 3.**
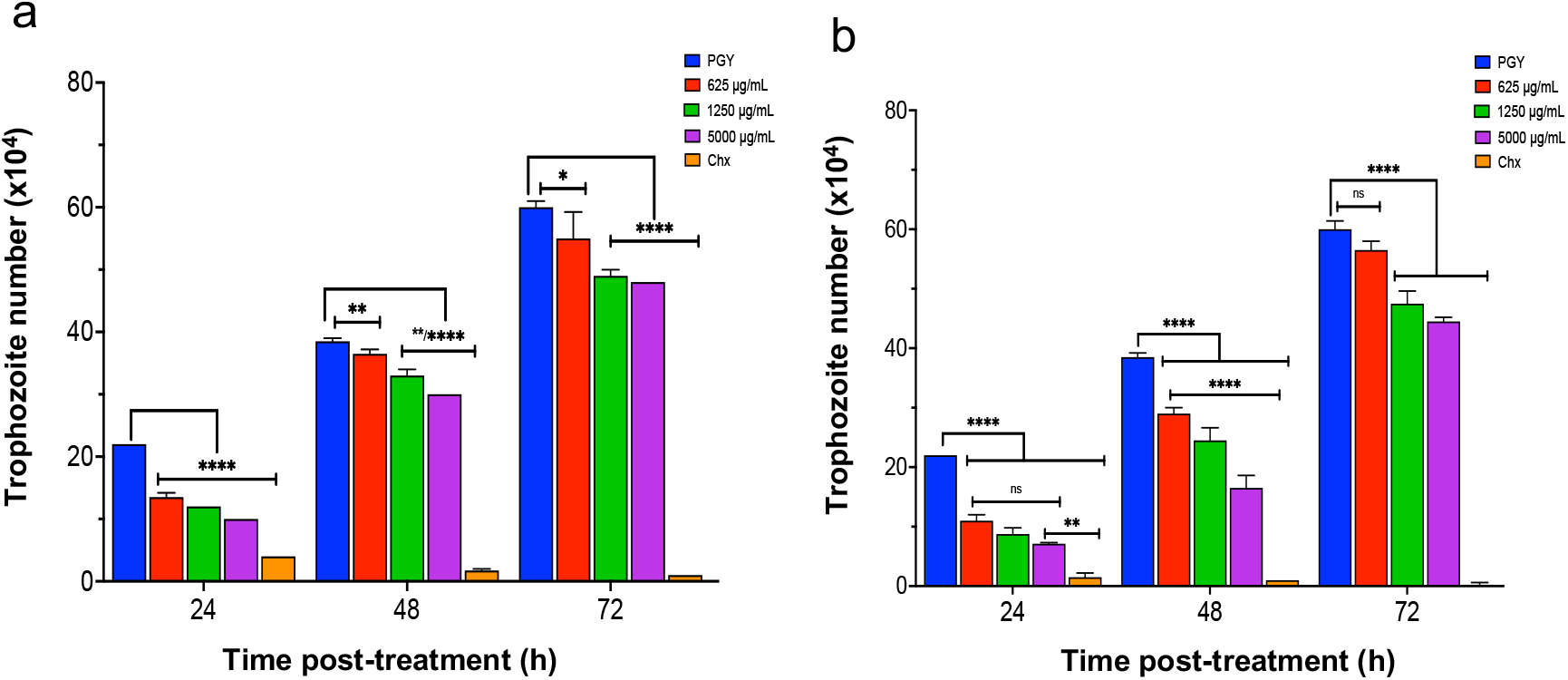
Transient effect of *Camellia* sinensis extract on trophozoites. Growth inhibition of trophozoites post-transient exposure to 625 µg/mL, 1250 µg/mL, and 5000 µg/mL of *C. sinensis* and Chx for 2 h (a) and 24 h (b). Significant differences were detected between Chx and the graded doses of *C*. sinensis at 72 h post-exposure at both time points. (** *P =* 0.0058; **** *P* ≤ 0.0001; ns, non-significant)

### 3.5. C. sinensis extract inhibits encystation

While determining the number of cysts at 72 h pre- and post-SDS digestion, One-way ANOVA revealed significant dose-dependent effects of *C. sinensis* on the encystation rate pre-SDS digestion (F (5, 12) = 5671, *P* < 0.0001) and post-SDS digestion (F (5, 12) =1909, *P* < 0.0001) (Fig. 4A). Post hoc comparisons of 2500 µg/mL and 5000 µg/mL *C. sinensis* showed no cysts with no significant difference with the positive control (5 mM PMSF) (*P* < 0.0001). The lower doses of 625 µg/mL and 1250 µg/mL showed some level of encystation but with significant difference with the negative control encystation buffer and the higher concentrations (*P* < 0.0001). One-way ANOVA revealed a significant main effect of *C. sinensis* (F (5, 12) =3149, *P* < 0.0001) on inhibition rate. Post hoc comparisons showed a 100% inhibition rate of 2500 µg/mL and 5000 µg/mL encystation buffer with no significant difference between them and the positive control (5 mM phenylmethylsulfonyl fluoride (PMSF)) (*P* < 0.0001). On the other hand, no significant difference was detected in encystation inhibition between lower concentrations 625 µg/mL and 1250 µg/mL, 30.67% and 35.57%, respectively with the negative control 29.10% (*p* > 0.05) (Fig. 4B).

**Fig. 4.**
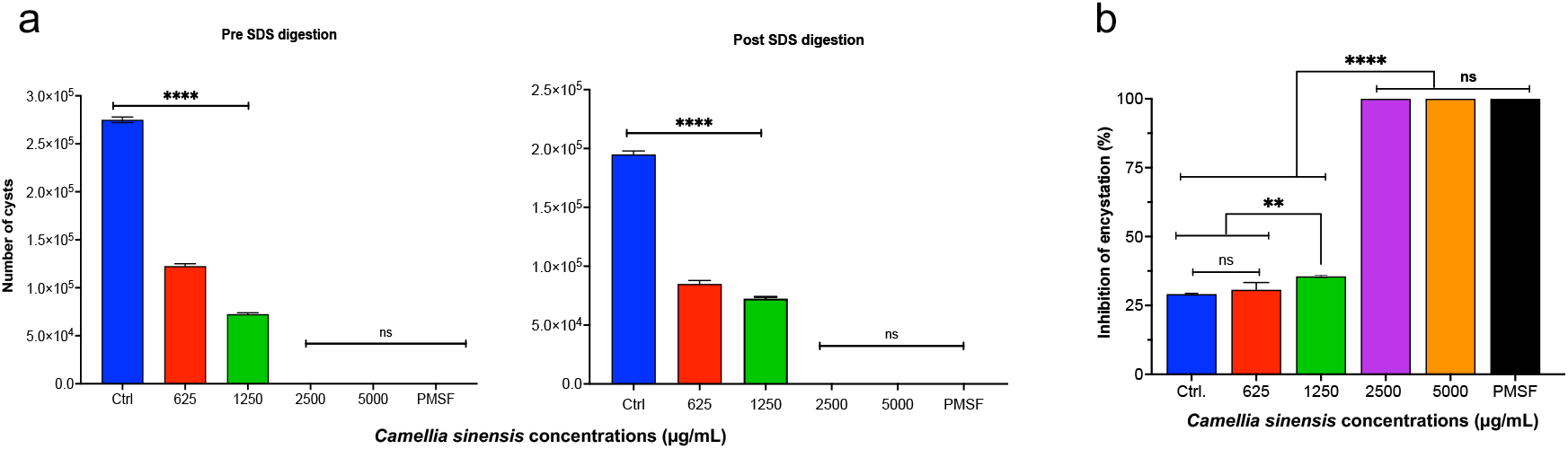
Effect of *Camellia sinensis* extract on encystation. (a) At 72 hours post synchronized encystation, pre- and post-SDS digestion, 2500µg/mL and 5000µg/mL *C. sinensis* showed not cysts with no significant difference with the positive control (5 mM PMSF). The lower doses of 625µg/mL and 1250µg/mL showed some level of encystation but with significant difference with the negative control encystation buffer and the higher concentrations. (b) Percentage inhibition of encystation of *A. castellanii*. 2500 µg/mL and 5000 µg/mL *C. sinensis* encystation buffer displayed 100% inhibition of encystation with no significant difference between it and the positive control (5 mM PMSF). The lower doses of 625 µg/mL and 1250 µg/mL show no significant difference with the negative control encystation buffer. (**** *P* < 0.0001; ns, non-significant)

### 3.6. C. sinensis extract interferes with cyst to trophozoite transformation

The effect of *C. sinensis* on *A. castellanii* cysts exposed to concentrations 312.5 -5000 *µ*g/mL was determined to check the cyst : trophozoite ratio at 72 h post exposure (Fig. 5A). One-way ANOVA revealed a significant main effect of *C. sinensis* concentrations (µg/mL) (F (7, 16) = 1001, *P* < 0.0001) on excystation rate of the cysts post-exposure. Post hoc comparisons showed that there was a significant dose-dependent decrease in excystation with 312.5-5000 *µ*g/mL *C. sinensis* (*P* < 0.0001), with an absolute inhibition of excystation at 1250 - 5000 *µ*g/mL (Fig. 5A). In determining the percentage inhibition of excystation, one-way ANOVA revealed a significant main effect of *C. sinensis* concentrations (µg/mL) (F (7, 16) =10695, *P* < 0.0001) on excystation inhibition percentage of the exposed cysts. Post hoc comparisons showed that 312.5 *µ*g/mL and 625 µg/mL concentrations exhibited 78% and 87% inhibition of excystation respectively with significant increase from the negative control groups (*P* < 0.0001). Similarly, the higher concentrations of 1250 - 5000 µg/mL displayed 100% inhibition of excystation mimicking the activity of CHX (*P* > 0.05) (Fig. 5B).

**Fig. 5.**
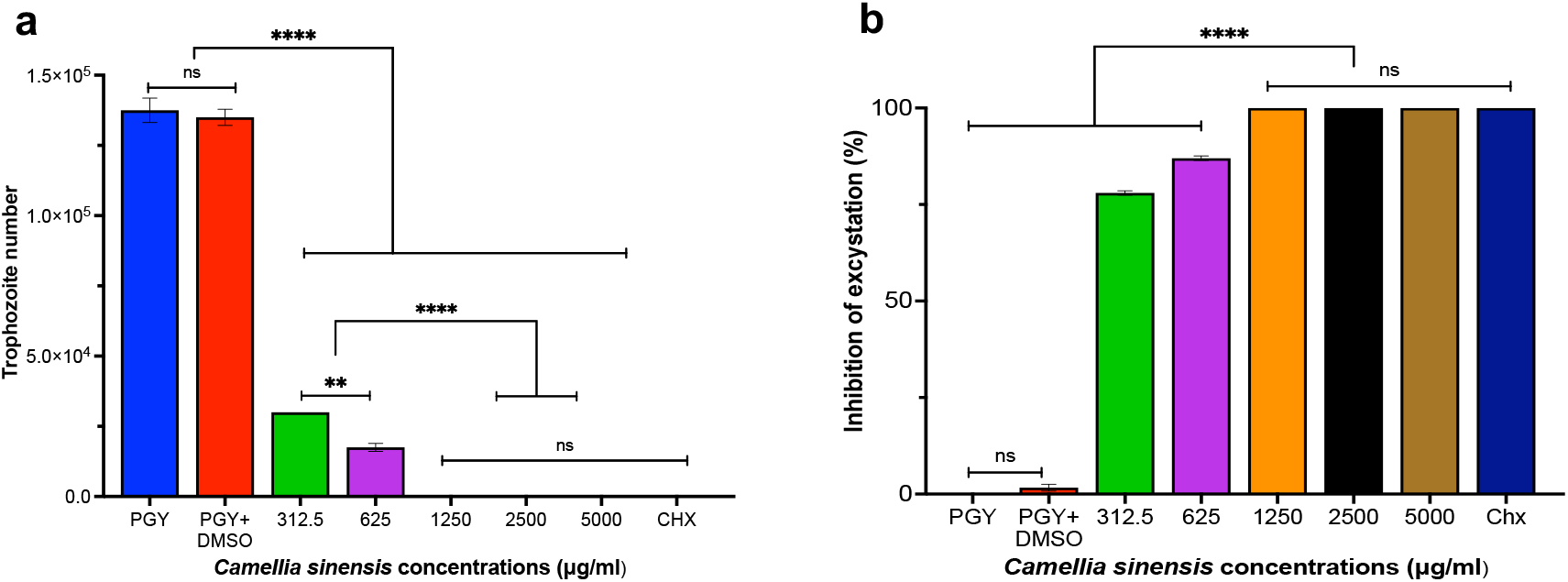
Effects of *C. sinensis* solvent extract on excystation. (a) 72 h post exposure to graded doses of *C. sinensis*, 1250 µg/mL, 2500 µg/mL and 5000µg/mL exhibited 100% inhibition of excystation as seen with Chx. Although there were significant differences between the higher doses and the lower doses of 312.5 µg/mL and 625 µg/mL, these low doses exhibited 78% and 87% inhibition of excystation respectively. (b) Percentage of trophozoite excystation in *C. sinensis* solvent extract: 72 h post exposure to graded doses of *C. sinensis*, 1250 µg/mL, 2500 µg/mL and 5000 µg/mL exhibited 100% inhibition of excystation as observed with Chx. Although there were significant differences between the higher doses and the lower doses of 312.5 µg/mL and 625 µg/mL, these low doses exhibited 78% and 87% inhibition of excystation, respectively. (**** *P* < 0.0001; ns, non-significant)

### 3.7. Morphological changes in A. castellanii after exposure to C. sinensis extract during excystation

Microscopic observation of cysts 72 h post exposure to *C. sinensis* showed extensive excystation of the negative control, while cysts exposed to *C. sinensis* exhibited mild to extensive damage as shown by the presence of damaged cysts, with cytosol contents in the culture media, suggesting cytolysis (Fig. S1). With increasing the concentrations of *C. sinensis*, the number of cellular debris increased. The debris observed with 312.5 *-* 625 µg/mL concentrations were not as extensive as those with the higher concentrations 1250 - 5000 µg/mL. To confirm the presence or absence of trophozoites, which adhere to the bottom of flasks, the flasks were shaken gently during microscopic imaging and the view of the entire flask bottom showed varying degree of floating debris and improper cysts with no sign of adherent trophozoites. Quantification of the number of cysts per aggregate was impossible to achieve as the gentle shaking seemed to loosen the inter-cystic adherent properties.

### 3.8. Ultrastructural features of active trophozoites and encysting trophozoites

SEM micrographs of trophozoites (Fig. 6) revealed rupture of plasma membrane, shrunken cytosol and changes in the size. Continued exposure to *C. sinensis* solvent extract from 24 h to 72h showed complete destruction of trophozoites. Chx showed similar morphological changes. The trophozoites in the control samples appeared encysting and cysts were amassed in bunches and held together by their adhesin. In contrast, exposure of encysting trophozoites to the solvent extract led to loss of their adhesive properties and breaking up of the cysts in a dose-dependent manner which was progressive even at 625 µg/mL. After 72 h exposure, there was extensive destruction of the encysting and encysted A. castellanii (Fig. 7).

**Fig. 6.**
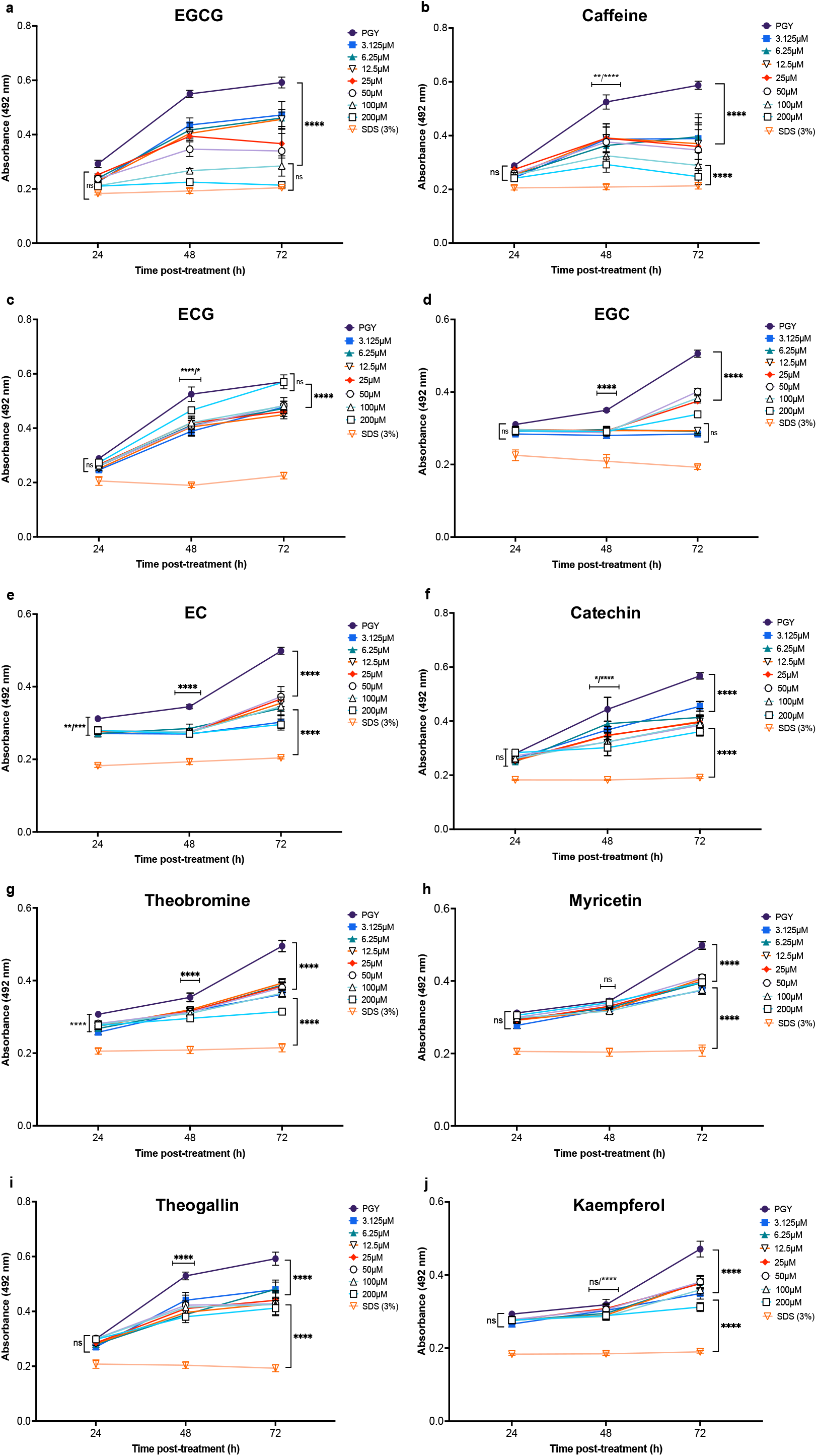
Growth inhibitory effect of *C. sinensis* constituents on *A. castellanii* at 24, 48, and 72 h post treatment with the indicated concentrations. (a) EGCG at 48 and 72 h post exposure, the PGY group showed significant increase in trophozoite viability, while at 72 h, 100 μM and 200 μM EGCG-treated groups showed no significant difference with the positive control (SDS) indicating that both groups successfully inhibited trophozoite replication. (b) Caffeine at 100 μM and 200 μM exhibited the same trend. (c-f). All treatment groups (ECG, EGC, EC, and catechin, respectively) showed significant difference to the negative control and to the positive control at 48 and 72 h post exposure. ECG and EC inhibited trophozoite replication at 48 h post treatment but were not able to sustain the effect and could not match the anti-acanthamoebic capability of positive control. (g-j) All treatment groups (theobromine, myricetin, theogallin, and kaempferol, respectively) showed significant difference to the negative control (PGY) and to the SDS at 48 and 72 h post exposure. Despite the differences these constituents did not exhibit sustained anti-acanthamoebic capability from 48 h post treatment as shown by SDS. (**** *P* ≤ 0.0001, ns: non-significant)

**Figure 7.**
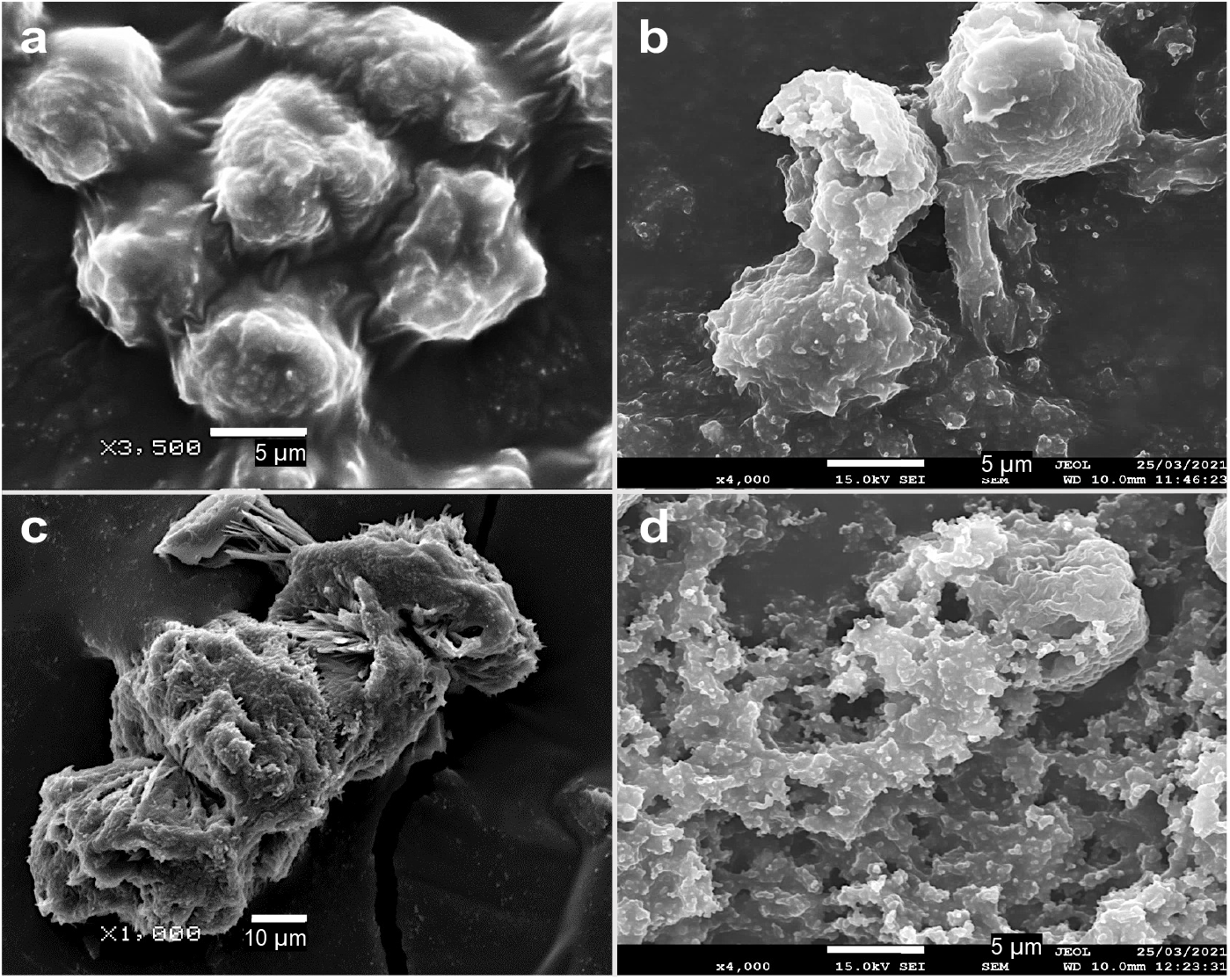
Representative scanning electron microscopy (SEM) micrographs of *A. castellanii* trophozoites showing loss of membrane of trophozoites and shrinking of their size possibly in the attempt to encyst. Continued exposure to both C. sinensis solvent extract and Chx showed loss of acanthopodia and progressive destruction of trophozoites. (a) Trophozoites in PGY (control) attached together by adhesin; (b) Trophozoites in 2500 µg/mL of C. sinensis solvent extract at 24 h showing loss of protective adhesin and progressive destruction of the trophozoites. (c) Trophozoites in Chx at 24 h showing loss of protective adhesin and shrinking/destruction of trophozoites. (d) Trophozoites in 2500 µg/mL of C. sinensis solvent extract at 72 h showing complete destruction of the trophozoites with some parasite debris in the background. Magnifications: X3500, X4000, X1000, and X4000 respectively. Scale bars, 5, 1,10, and 1 µm, respectively.

### 3.9. Chromatographic profile of C. sinensis solvent extract

The first mobile phase (acetic acid + acetonitrile) revealed 6 fractions (Fig. S2), while from the second mobile phase (0.1% formic acid in methanol) (Fig. S3) 10 fractions were identified; fractions are represented as peaks at different retention times. The chromatograph with mobile phase of 0.1% formic acid in water and water + 0.1% formic acid in methanol showed more individual peaks and another peak which showed coelution of two analytes at 1-hour retention time. To identify the analytes in the fractions observed in the chromatographs, samples of the fractions were characterized with high-performance liquid chromatography (HPLC).

### 3.10. Liquid Chromatography-Mass Spectrometry (LC-MS)

From the 10 fractions separated during the flash column chromatography of *C. sinensis* solvent extract, fraction 1 did not contain any components as the molecular weights detected could not be identified against any database. The nine remaining fractions showed 10 individual components (Table 1) with the 7^th^ and 8^th^ component showing as a coelution of two compounds in one fraction thereby making it a total of 10 components in the fractions. The coelution occurred due to similarity in their monomeric isotope mass and molar mass.

**Table 1.**
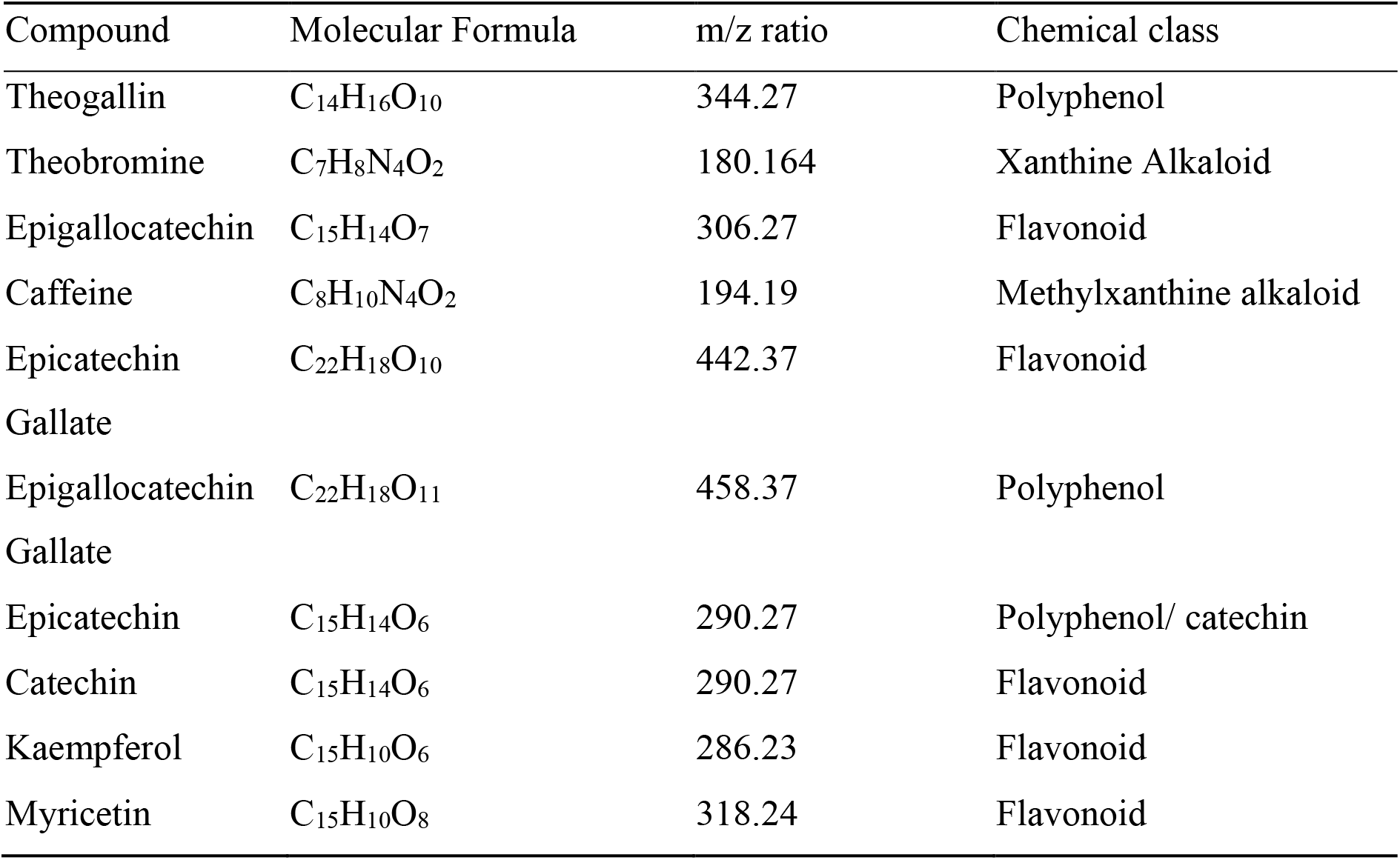
Chemical constituents of *C. sinensis* solvent extract analysed by UHPLC–QTOF–MS.

### 3.11. Trophocidal effects of C. sinensis components

The growth and replicatory inhibitory effect of *C. sinensis* ingredients identified by UHPLC–QTOF–MS analysis (Table 1) on trophozoites at doses of 3.125 µM, 6.25 µM, 12.5 µM, 25 µM, 50 µM, 100 µM and 200 µM at 24, 48, and 72 h post treatment. The results of individual standards are as follows.

For EGCG, at 24 h post- treatment, post hoc comparisons of treatment groups with both controls (negative and positive control (SDS 3%)) showed that only the negative control group exhibited any form of significant trophozoite growth while all the treatment groups showed no growth with no significant difference in their trophozoite quantity when compared to the positive control (*P* > 0.05). This trend however was not sustained for the 48 and 72 h post-treatment comparisons as 100 µM and 200 µM concentrations were the only doses with similar growth inhibitory capabilities of the SDS control groups. The lesser concentrations showed continuous trophozoite growth though at a lesser rate than the negative control group. The 25 µM and 50 µM concentrations showed a halt and decline of trophozoite growth after 48 h post-treatment, but the efficacy had significant difference between those of 100 µM and 200 µM which mirrored the effect of SDS control and eliminated *A. castellanii* trophozoites with no statistical difference (*P* > 0.05) (Fig. 8A).

**Figure 8.**
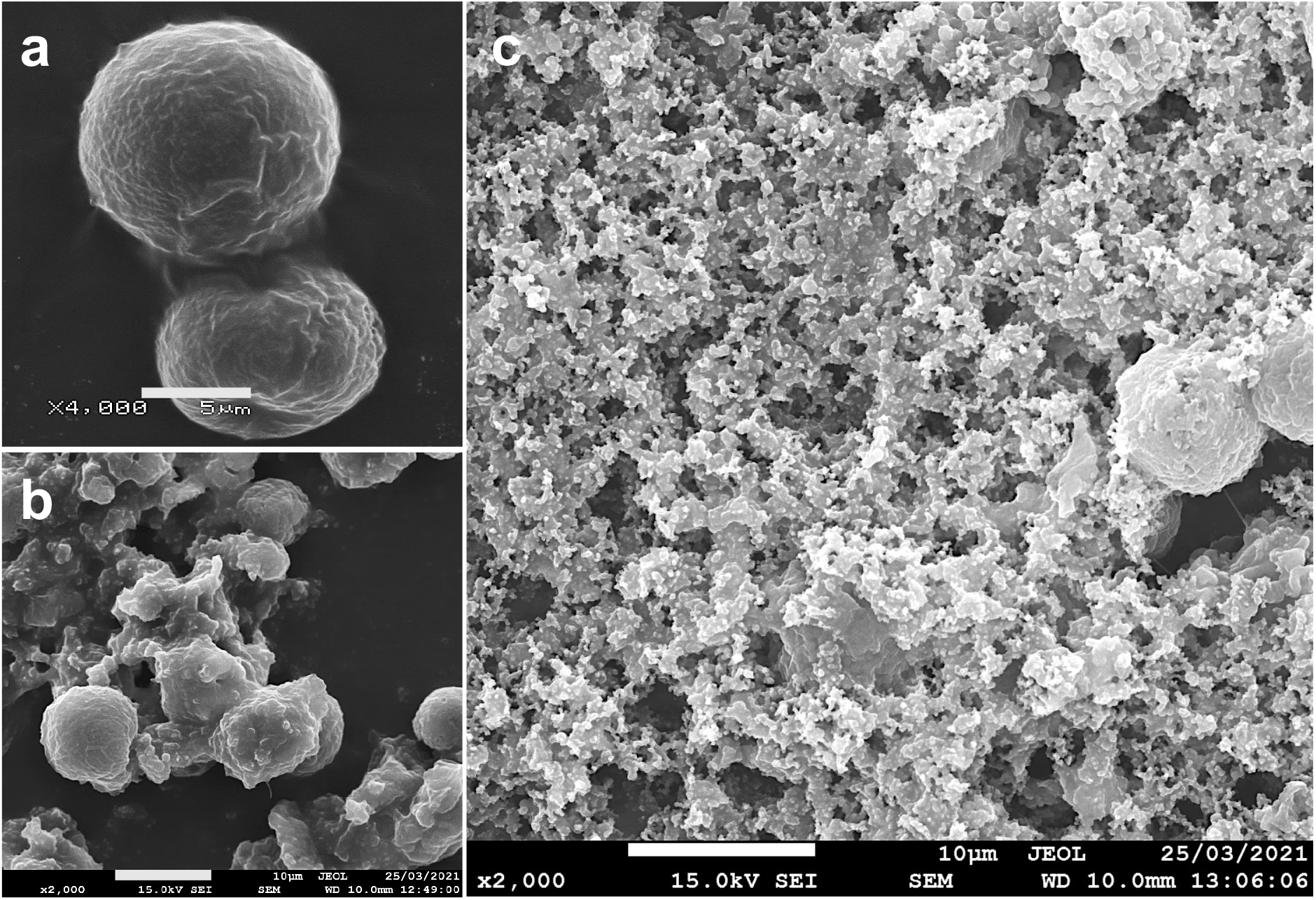
Representative scanning electron microscopy (SEM) micrographs showing destruction of *A. castellanii* cysts exposed *to C. sinensis* solvent extract. **(a)** Cysts in the control encystation buffer adhered together; (b) Aggregated cysts exposed to *C. sinensis* 625µg/mL at 72 h showing a number of broken cysts, cyst debris and a few intact cysts; (c) Cysts exposed to *C. sinensis* 2500 µg/mL for 72 h showed extensive destruction. Magnifications: X4,000, X2,000, and X4,000, respectively; Scale bars, 5, 10, and 10 µm, respectively.

For caffeine, at 24 h post- treatment, post hoc comparisons of treatment groups with both controls (negative control, positive control (SDS 3%)) showed that only the negative control group exhibited any form of significant trophozoite growth while all the treatment groups showed none with no significant difference in their trophozoite quantity when compared to the SDS control group (*P* > 0.05). As seen with EGCG, this trend however was not sustained for the 48h post-treatment comparisons as all the concentrations showed growth of trophozoites though with significance difference from the PGY control growth rate. Interestingly after 48 h, all the treatment concentrations except for 3.125 µM and 6.25 µM exhibited halt of and decline of trophozoite growth with the 100 µM and 200 µM concentrations mirrored the effect of SDS control at 72 h post-treatment and eliminated *A. castellanii* trophozoites with no statistical difference (*P* > 0.05) (Fig. 8B).

For the other catechins, the two-way RM ANOVA of ECG revealed a significant main effect of time (h) (F (2, 6) = 104.6, *p* < 0.0001, concentration (µg/ml) (F (8, 24) =91.71, *P* < 0.0001), and time x concentration interaction (F (16, 48) = 11.86, *P* < 0.0001). While for EGC, the two-way RM ANOVA revealed a significant main effect of time (h) (F (2, 6) = 152, *P* = 0.0002, concentration(µg/mL) (F (8, 16) =158.1, *P* < 0.0001), and time x concentration interaction (F (16, 32) = 63.21, *P* < 0.0001). For EC, the two-way RM ANOVA revealed a significant main effect of time (h) (F (2, 6) = 40.45, *P* = 0.0003, concentration (µg/mL) (F (8, 24) =54.61, *P* < 0.0001), and time x concentration interaction (F (16, 48) = 22.95, *P* < 0.0001). For Catechin, the two-way RM ANOVA revealed a significant main effect of time (h) (F (2, 4) = 14.65, *P* = 0.0144, concentration(µg/mL) (F (8, 16) = 46.69, *P* < 0.0001), and time x concentration interaction (F (16, 32) = 8.277, *P* < 0.0001). Post hoc comparisons of catechin treatment groups with both controls (negative control, positive control (3% SDS) showed that although there was continuous significance difference between the negative control group and the catechins, they all mostly exhibited little or no form of inhibition to the trophozoite growth from 24 h to 72 h post treatment to all the doses of *C. sinensis* components (Fig. 8C-F). ECG was the only catechin that showed any notable inhibition to trophozoite growth as a selective range of low concentrations of 3.125 µM, 6.25 µM and 12.5 µM. Although these concentrations did not destroy the trophozoites, they interestingly inhibited the trophozoite replication in sort suspended animation resulting neither in the replication or the depletion of the OD recorded for the concentrations (Fig. 8C).

For the remaining *C. sinensis* components, the two-way RM ANOVA of theobromine revealed a significant main effect of time (h) (F (2, 6) = 952.2, *P* < 0.0001, concentration(µg/mL) (F (8, 24) = 223.7, *P* < 0.0001), and time x concentration interaction (F (16, 48) = 52.50, *P* < 0.0001). While for myricetin, the two-way RM ANOVA revealed a significant main effect of time (h) (F (2, 6) = 2224, *P* = 0.0002, concentration(µg/ml) (F (8, 24) =348.7, *P* < 0.0001), and time x concentration interaction (F (16, 48) = 32.99, *P* < 0.0001). For theogallin, the two-way RM ANOVA revealed a significant main effect of time (h) (F (2, 6) = 20.47, *P* = 0.0021, concentration (µg/ml) (F (8, 24) = 111.9, *P* < 0.0001), and time x concentration interaction (F (16, 48) = 15.86, *P* < 0.0001). For kaempferol, the two-way RM ANOVA revealed a significant main effect of time (h) (F (2, 6) = 90.28, *P* < 0.0001, concentration(*µ*g/mL) (F (8, 24) = 97.90, *P* < 0.0001), and time x concentration interaction (F (16, 48) = 46.7, *P* < 0.0001). Post hoc comparisons of the analysed treatment groups with both controls (negative control, positive control (SDS 3%) showed that although there was interchangeable continuous significance difference between the negative control group and the catechins, they all exhibited no inhibition to the trophozoite growth from 24 h to 72 h post treatment to all the doses of *C. sinensis* components (Fig. 8G-J).

An analysis of the all the tested *C. sinensis* components combined in a full complement of a 1:1 ratio for all the chemicals was done. A two-way RM ANOVA of the components complement revealed a significant main effect of time (h) (F (2, 8) = 420, *P* < 0.0001, concentration (µg/mL) (F (8, 32) = 99.29, *P* < 0.0001), and time x concentration interaction (F (16, 48) = 12.47, *P* < 0.0001). Post hoc comparisons of the analysed complement with both controls (negative control, positive control (SDS 3%) showed that despite the individually ranged efficacies of the chemicals making up the complement, there seemed to be no synergistic activity of these chemicals as the complement exhibited no inhibition to trophozoite growth from 24 h to 72 h post treatment to all concentrations of *C. sinensis* ingredients (Fig. 9). EGCG and caffeine were the only *C. sinensis* constituents that inhibited trophozoite replication within the safe concentration of 3.125 µM to 200 µM, with no significant difference with the efficacy exhibited by SDS control.

**Fig. 9.**
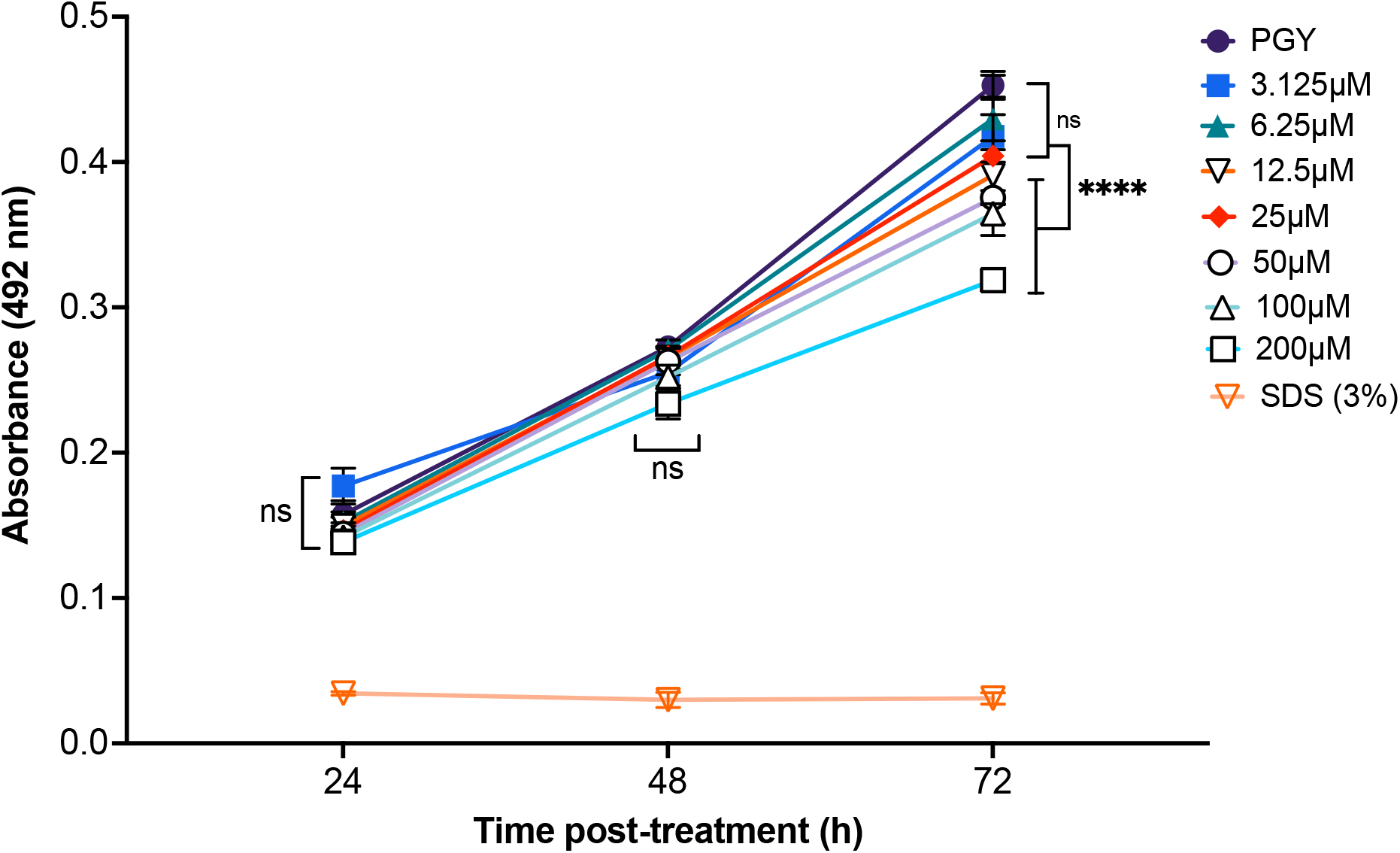
Growth inhibitory effect of *C. sinensis* components combined at equal proportions. The combined treatment included EGCG, ECG, EGC, EC, Catechin, Caffeine, Theobromine, Myricetin, Theogallin, and Kaempferol. Trophozoites were treated with 3.125 µM, 6.25 µM, 12.5 µM, 25 µM, 50 µM, 100 µM, and 200 µM for 24, 48, and 72 h. No significant differences were detected between treated groups and negative control (PGY) at 24 h and 48 h, while only 50 µM, 100 µM and 200 µM showed significant difference compared to PGY treatment. Despite the differences, the chemical constituents did not exhibit anti-acanthamoebic capability for the duration of the experiment with significant difference between them and positive control (SDS). (**** *P* ≤ 0.0001, ns: non-significant)

### 3.12. Cysticidal effect of C. sinensis ingredients

One-way ANOVA revealed a significant main effect of EGCG (F (5, 12) =3606, *P* < 0.0001), theobromine (F (5, 12) =3606, *P* < 0.0001), and EGCG + theobromine (F (5, 12)=3606, *P* < 0.0001) on the encystation inhibition rate. Interestingly, post hoc comparisons showed a 100% trophozoite encystation inhibition rate of all tested concentrations of *C. sinensis* chemical ingredients in hyperosmotic solutions, with no significant difference between them and the positive control (5 mM PMSF) (*P* < 0.0001) (Fig. 10).

**Fig. 10.**
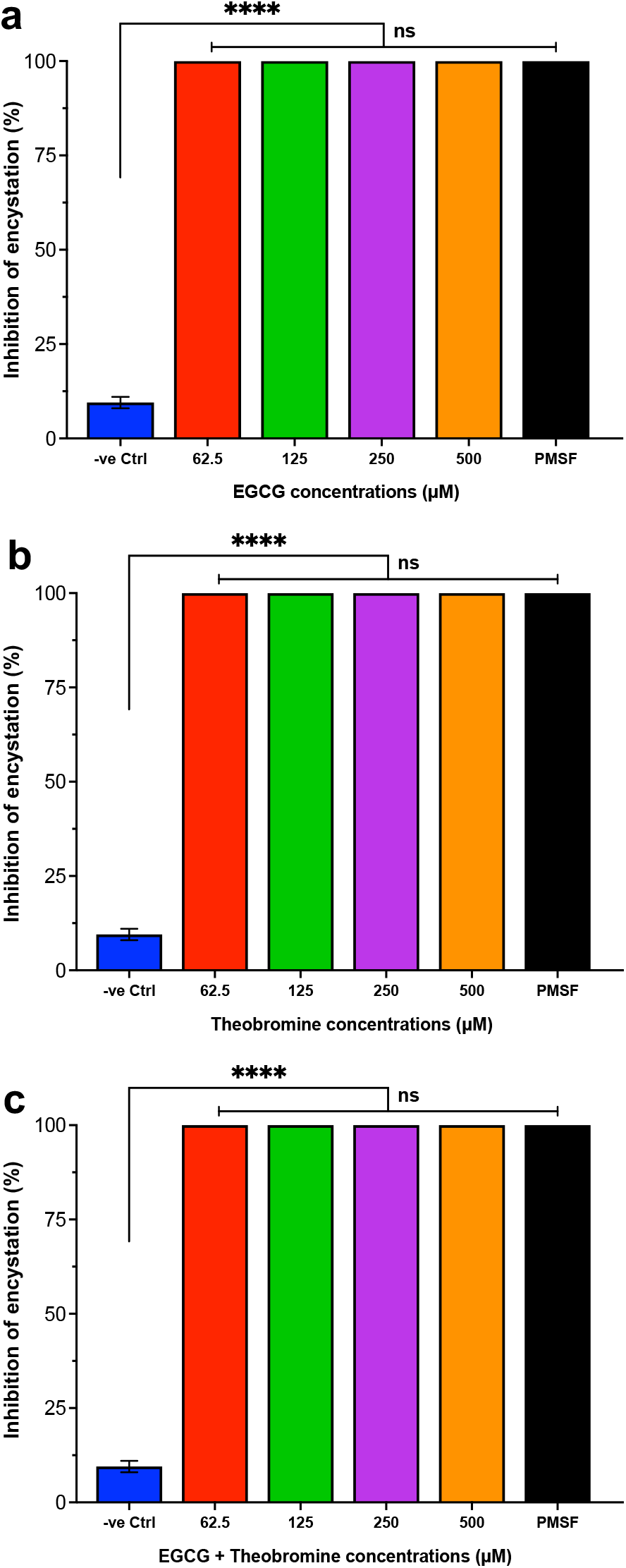
Inhibition of encystation of *A. castellanii* exposed to two *Camellia sinensis* components individually (a: EGCG, b: Theobromine) and as a combination at a 1:1 ratio (c). EGCG and Theobromine encystation solutions were used at concentrations of 62.5 µM, 125 µM, 250 µM, and 500 µM. All concentrations displayed 100% inhibition of encystation with no significant difference between them and positive control (5 mM PMSF). (**** *P* ≤ 0.0001, ns: non-significant)

## 4. Discussion

The limitations of current therapies of *Acanthamoeba* keratitis warrants the development of improved treatment approaches. In this regard, *C. sinensis* brews have shown great promise, exhibiting trophocidal and cysticidal activities (Fakae et al., 2020). Herein, we examined the efficacy of *C. sinensis* solvent extract as an anti-acanthamoebic agent and identified and tested the anti-acanthamoebic efficacy of individual components of the solvent extract.

The present study evaluates the toxicity of *C. sinensis* solvent extract on HCE-2 cells and MDCK cells *in vitro*. Our results demonstrate that *C. sinensis* exhibits dose-dependent toxic effect on MDCK cells, particularly at concentrations 1250-5000 µg/mL. In contrast, HCE-2 cells, exposed to 2500 µg/mL, exhibited an initial 7.5% cytotoxic effect after 24 h, which was dropped to ∼5% at 48 h. Even the highest concentration of 5000 µg/mL exhibited only 30% cytotoxicity for HCE-2 cells up to 48 h post exposure. Selectivity index showed that *C. sinensis* solvent extract has selective cytotoxicity for *A. castellanii* as observed by the wide range of concentration allowed for HCE-2 cells. However, MDCK cells were more sensitive to toxicity caused by *C. sinensis* than trophozoites. The choice of continuing the experiment with the range of concentrations of *C. sinensis* became dependent on the cytotoxicity level observed in the HCE-2 cells, given the potential use of *C. sinensis* as a topical agent against AK, which has the cornea as its predilection site (Siddiqui and Khan, 2012; Lorenzo-Morales et al., 2013).

We demonstrated the anti-acanthamoebic activity of *C. sinensis* solvent extract against trophozoites and cysts. The concentrations 625*-*5000 µg/mL exhibited a sustained and progressive inhibition of trophozoite replication and cytolysis. The lower concentrations of *C. sinensis* solvent extract 625*-*1250 µg/mL slightly inhibited encystation, while higher concentrations, 2500*-*5000 µg/mL, completely inhibited trophozoite-to-cyst differentiation. In addition to inhibition of encystation, *C. sinensis* caused a destruction of trophozoites as they attempt to encyst. Electron microscopic analysis showed that exposure of *A. castellanii* to *C. sinensis* solvent extract resulted in considerable signs of cellular damage, including rupture of plasma membrane and disruption of the adhesin. In evaluating the effect of *C. sinensis* solvent extract on the differentiation of *A. castellanii* from cysts to trophozoites, lower concentrations partially and higher concentrations fully inhibited excystation. Light microscopic analysis of the culture at 72 h post exposure showed extensive destruction of *A. castellanii*. It is intriguing that trophozoites exposed to the same low concentration 312.5 µg/mL of *C. sinensis* solvent extract which did not inhibit trophozoite replication, was able to inhibit excystation and destroyed cysts and trophozoites during excystation. Although *A. castellanii* can encyst when it is exposed to adverse conditions, trophozoites might not had the ability to halt and reverse its excystation process in the presence of *C. sinensis* in the culture media. This disruption in the phenotypic differentiation of *A. castellanii* may have caused an activation of a proapoptotic pathway and death of the excysting trophozoites.

A previous chromatographic analysis of *C. sinensis* brew identified caffeine, ECG, EGCG, theogallin, quercetin, and kaempferol. In the present study, five additional compounds – Theobromine, EGC, EC, catechin, and myricetin – were identified in the chromatographic analysis of the solvent extract. It is unclear why the anti-acanthamoebic activity of a pooled mix of the identified compounds was limited. It would seem, however, that the combinations of all or some of the chemical compounds of *C. sinensis* outside its natural constitution may not have yielded desirable results because a host of other constituents of cellulose, proteins, amino acids and other compounds in the solvent extract may not have been accounted for. Although a combination of *C. sinensis* bioactive compounds did not yield a favourable anti-acanthamoebic activity, some of these compounds exhibited a sustained amoebicidal activity. The evaluation of individual bioactive compounds showed that EGCG and caffeine exhibited a dose-dependent trophocidal activity at 72 h post-exposure.

The combination of EGCG with *C. sinensis* matcha tea displayed anti-acanthamoebic activity against trophozoites and cysts (Dickson et al., 2020). EGCG has shown antifolate activity against *Stenotrophomonas maltophilia* (Navarro-Martínez et al., 2005), via inhibiting dihydrofolate reductase (DHFR), a key enzyme that catalyses the reduction of 7,8-dihydrofolate to 5,6,7,8-tetrahydrofolate, which is involved in nucleotide biosynthesis. Interestingly, *A. castellanii* has DHFR and the antifolate trimethoprim drug has amoebicidal activity (Siddiqui et al., 2016). Thus, inhibition of DHFR by EGCG can disrupt DNA synthesis in *A. castellanii*. EGCG can inhibit protein synthesis and induce cellular apoptosis in pancreatic cancer cells (Shankar et al., 2007). Additionally, EGCG has a broad-spectrum inhibitory activity against virus attachment to host cells by regulating protein synthesis of host cell (Ciesek et al., 2011; Colpitts et al., 2014). It is therefore conceivable that EGCG, via its antifolate and protein synthesis inhibitory activities, may contribute to the anti-acanthamoebic effects of *C. sinensis*.

*A. castellanii* secretes many serine protease and metalloprotease (Khan, 2006), which compromise cell membrane integrity and cause cytolysis. In addition to being determinants of the protozoan virulence (Dudley et al., 2008), these enzymes can also regulate *A. castellanii* trophozoite into cyst differentiation (Lorenzo-Morales et al., 2015; Alsam et al., 2005). A study of *C. sinensis* polyphenols showed that EGCG appears to be a natural inhibitor of serine protease and metalloprotease (Benelli et al., 2002; Wyganowska-Świątkowska et al., 2018). This inhibitory capability seems to be broad spectrum as it also inhibits 3 chymotrypsin-like protease (3CLpro) of SARS-CoV-2, resulting in inhibition of viral polyprotein processing and presumably inhibiting spread of coronavirus (Chiou et al., 2021). EGCG can increase the expression of tissue factor pathway inhibitor-2 (TFPI-2), a Kunitz-type serine proteinase inhibitor (Gu et al., 2009). Therefore, *C. sinensis*-derived EGCG, via the interaction with or inhibition of these enzymes, might have caused cytolysis of *A. castellanii* trophozoites and inhibited encystation.

Caffeine also inhibited trophozoite replication. Caffeine can regulate the activation of cAMP synthesis, which regulates chemotaxis and gene transcription in the amoeba *Dictyostelium discoideum* (Brenner and Thoms, 1984). There are similarities in the cAMP-specific phosphodiesterase, RegA, in *A. castellanii* and *D. discoideum*, which plays a role in regulating spore formation in *D. discoideum* and inhibiting encystation in *A. castellanii by increasing cAMP levels (Du et al*., *2014). It is therefore possible that the dose-dependent destruction of trophozoites 72 h post-exposure to caffeine might be attributed to increased cAMP levels which may have interfered with encystation. Contrary to the inability to inhibit trophozoite replication as exhibited by caffeine and EGCG, interestingly, theobromine was able to inhibit cyst to trophozoite differentiation during excystation even at the lowest concentration of* 62.5 µM. *Also, a combination of theobromine and EGCG inhibited trophozoites’ encystation*.

## 5. Conclusion

*C. sinensis* solvent extract had amoebicidal activities against both trophozoite and cyst forms of *A. castellanii*. While *C. sinensis* solvent extract exhibited dose-dependent amoebicidal ability, a combination of chemical components of *C. sinensis* did not. However, *C. sinensis* components with the most amoebicidal ability (EGCG, Caffeine and Theobromine) exhibited 100% inhibition of trophozoite encystation. Additionally, *EGCG and caffeine* had amoebicidal ability against both forms of *A. castellanii*, and catechins and theobromine had cysticidal effects. Knowledge gleaned from this study provide insight into the anti-acanthamoebic activity of *C. sinensis*, which may help in the development of novel topical therapeutics against AK. Future studies should investigate the *in vivo* toxicity and anti-acanthamoebic efficacy of *C. sinensis* and to elucidate the detailed molecular mechanisms leading to the damage of *A. castellanii* after exposure to *C. sinensis*.

## Credit authorship contribution statement

**Lenu B. Fakae:** Methodology, investigation, data curation and original draft preparation; **Mohammad S. R. Harun:** Software and data curation; **Darren Shu Jeng Ting:** Reviewing and investigation; **Harminder S. Dua:** Reviewing; **Gareth W.V. Cave:** Methodology, data curation and software; **Xing-Quan Zhu:** Validation and reviewing; **Carl W. Stevenson** Supervision, validation, and reviewing; **Hany M. Elsheikha:** Conceptualization, validation, visualisation, supervision and reviewing.

## Data availability

All data are provided in the article.

## Acknowledgments

We thank the University of Nottingham School of Life Sciences imaging unit (SLIM) for technical assistance.

## Appendix A. Supplementary data

Supplementary data to this article can be found online at

## Supplementary data

**Fig. S1.**
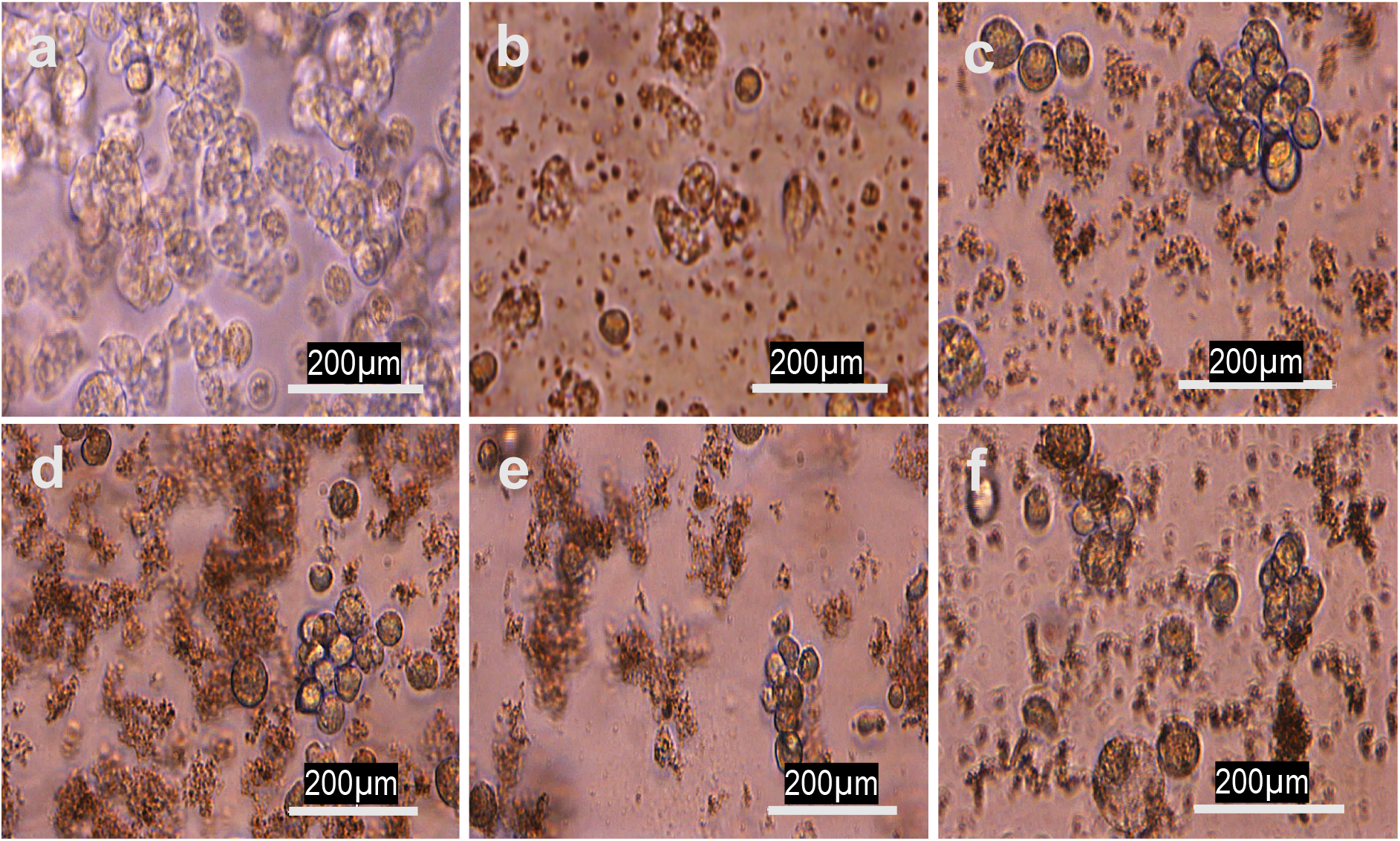
Representative photographs of *A. castellanii* cysts at 72 h post exposure to 312.5 – 5000 µg/mL *C. sinensis* extract using phase-contrast microscopy. **(a)** Control cyst culture treated with PGY. **(b-f)** Cysts treated with 312.5 µg/mL, 625 µg/mL, 1250 µg/mL, 2500 µg/mL and 5000 µg/mL *C. sinensis*, respectively. **(d-f)** 1250 µg/mL, 2500 µg/mL and 5000 µg/mL *C. sinensis* showed the presence of damaged cysts, with cytosol contents observed in the culture medium. Adhesion of trophozoites to flask surface was inhibited by all *C. sinensis* concentrations confirmed by the gentle rocking of the culture plates. **(b, c)** Cultures treated with 312.5 µg/mL and 625 µg/mL, respectively also showed some debris and damaged cysts but not as extensive as observed in the higher concentrations. X40 magnification. Scale bar, 200 µm.

**Fig. S2.**
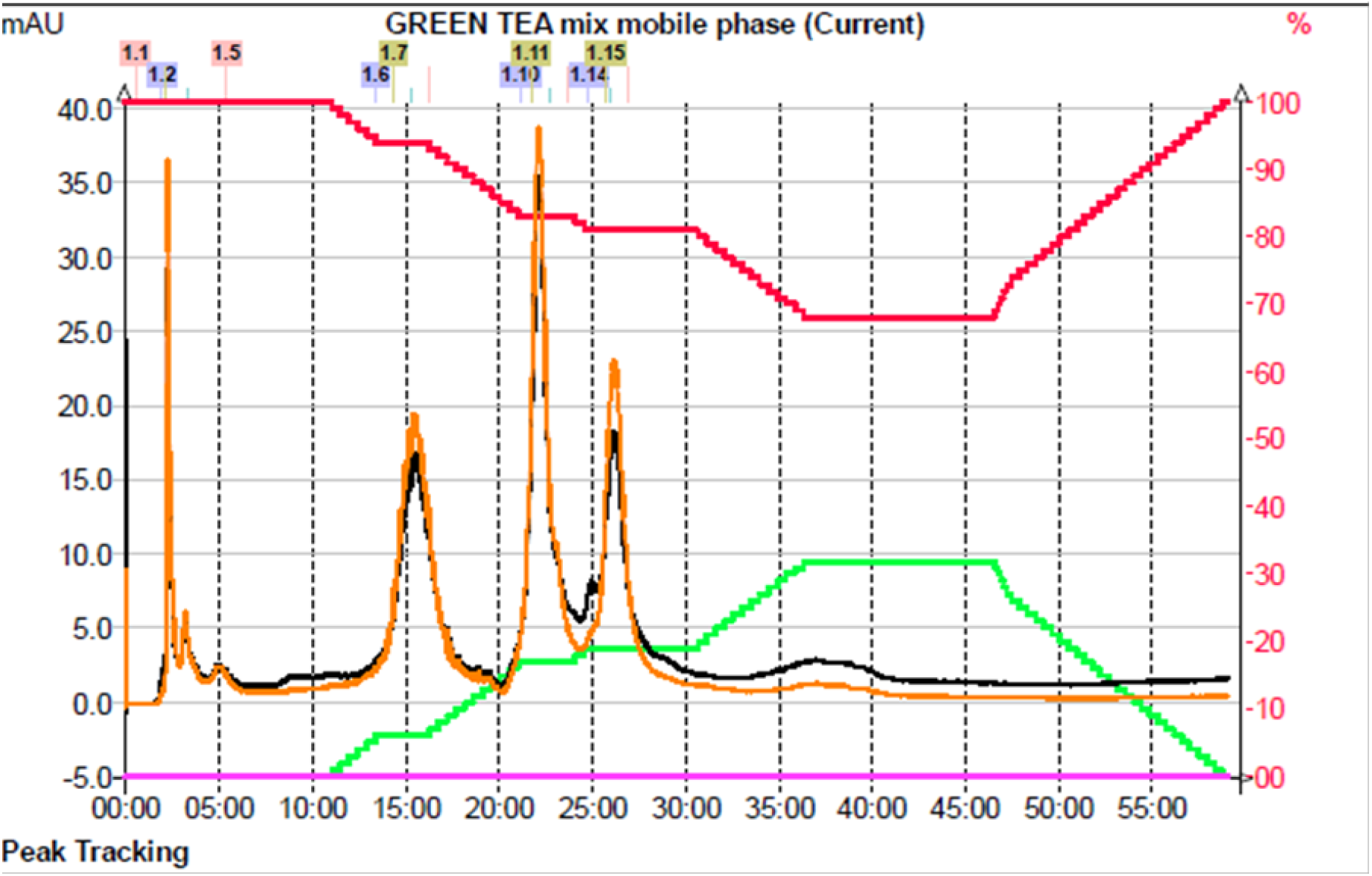
Flash column chromatography of *C. sinensis* solvent extract, mobile phase 1. *C. sinensis* solvent extract Flash chromatograph with mobile phase 0.2 % acetic acid in water + acetonitrile at a 91:9 ratio and water + acetonitrile at a 20:80 ratio showing peaks representative of fractions with possible analytes present in *C. sinensis* solvent extract.

**Fig. S3.**
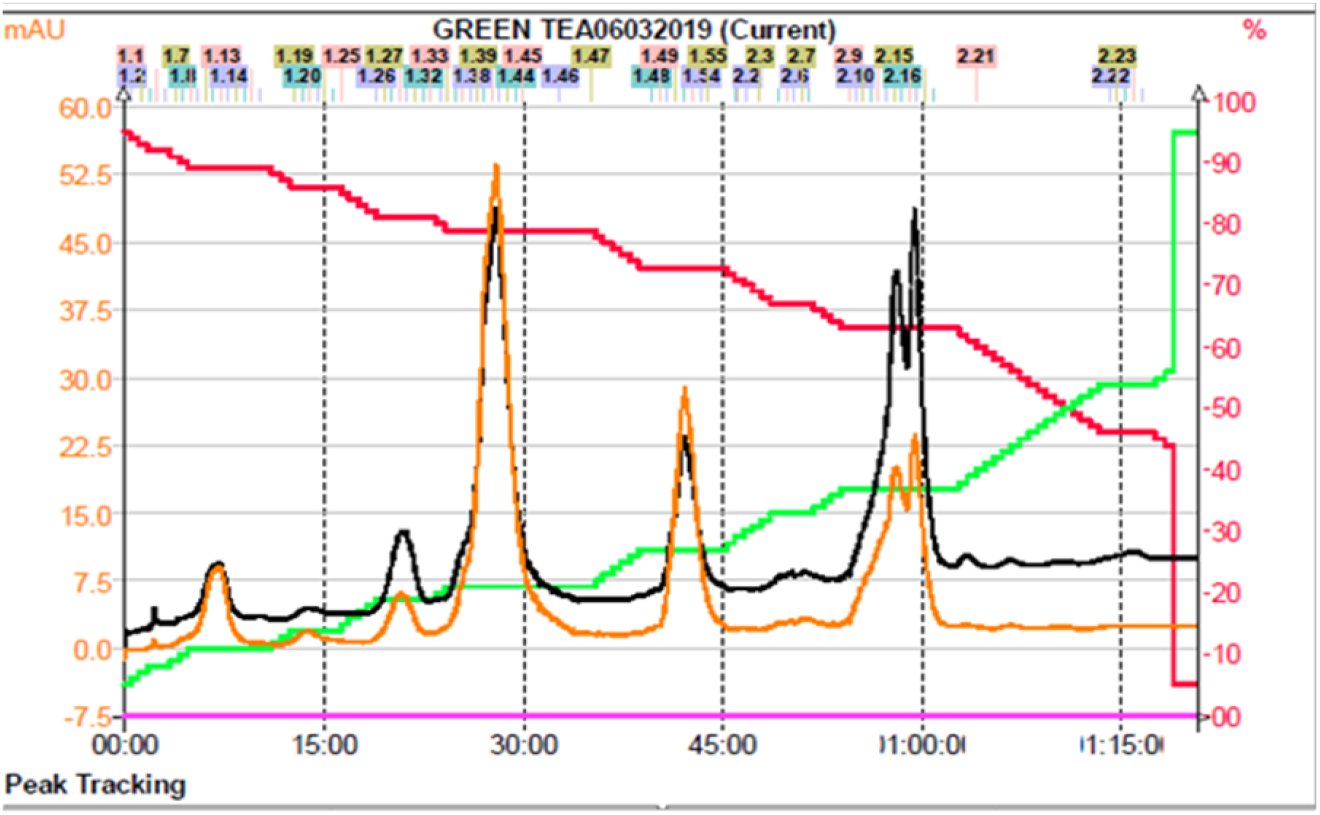
Flash column chromatography of *C. sinensis* solvent extract, mobile phase 2. *C. sinensis* solvent extract Flash chromatograph with mobile 0.1% formic acid in water and water + 0.1% formic acid in methanol showing peaks representative of fractions with possible analytes present in *C. sinensis* solvent extract.

